# Joint Gene Network Construction by Single-Cell RNA Sequencing Data

**DOI:** 10.1101/2021.07.14.452387

**Authors:** Meichen Dong, Yiping He, Yuchao Jiang, Fei Zou

## Abstract

In contrast to differential gene expression analysis at single gene level, gene regulatory networks (GRN) analysis depicts complex transcriptomic interactions among genes for better understandings of underlying genetic architectures of human diseases and traits. Recently, single-cell RNA sequencing (scRNA-seq) data has started to be used for constructing GRNs at a much finer resolution than bulk RNA-seq data and microarray data. However, scRNA-seq data are inherently sparse which hinders direct application of the popular Gaussian graphical models (GGMs). Furthermore, most existing approaches for constructing GRNs with scRNA-seq data only consider gene networks under one condition. To better understand GRNs under different but related conditions with single-cell resolution, we propose to construct Joint Gene Networks with scRNA-seq data (JGNsc) using the GGMs framework. To facilitate the use of GGMs, JGNsc first proposes a hybrid imputation procedure that combines a Bayesian zero-inflated Poisson (ZIP) model with an iterative low-rank matrix completion step to efficiently impute zero-inflated counts resulted from technical artifacts. JGNsc then transforms the imputed data via a nonparanormal transformation, based on which joint GGMs are constructed. We demonstrate JGNsc and assess its performance using synthetic data. The application of JGNsc on two cancer clinical studies of medulloblastoma and glioblastoma identifies novel findings in addition to confirming well-known biological results.

## 1. Introduction

Complex transcriptomic interactions exist among genes (Allocco et al., 2004; Trapnell et al., 2014). Deciphering such genetic architecture is essential in understanding the mechanism of complex diseases (Buil et al., 2015). Gene regulatory network (GRN) analysis is an attractive way to build transcriptional relationships among genes in providing a comprehensive view of gene expression dependencies (Wolfe et al., 2005; Peng et al., 2005; Margolin et al., 2006; Faith et al., 2007; Langfelder and Horvath, 2008; Sales and Romualdi, 2011; Ballouz et al., 2015; Ha et al., 2015; van Dam et al., 2018; Grimes et al., 2019; Ramirez et al., 2020; Van de Sande et al., 2020). In a GRN, individual genes are represented as nodes and gene pairs with interactions are connected by edges (Chen and Mar, 2018).

In practice, one focus area in biological and clinical research is the study of GRN changes across different conditions since transcriptomic interactions often vary across different tissues, cell types and cell states. Estimating a single network that applies to multiple conditions may not be appropriate and can lead to a biased estimation. In contrast, separately constructing graphical models under each condition is statistically valid (Lee et al., 2018). However, its performance may be improved by utilizing the network similarity across the related conditions. As such, joint graphical modeling is preferred (Danaher et al., 2014; Oates et al., 2014; Chun et al., 2015; Yang et al., 2015; Haslbeck and Waldorp, 2015; Cai et al., 2016; Qiu et al., 2016; Hallac et al., 2017; Huang et al., 2017; Lyu et al., 2018; Zhu and Koyejo, 2018; Lee et al., 2018; Geng et al., 2019; Jia and Liang, 2019; Zhang et al., 2019; Wu et al., 2019, 2020).

GGMs and their adaptive models are popular for constructing GRNs with microarray data due to their ease of interpretation. Recently, bulk RNA-seq data and newly emerging scRNA-seq data enable researchers to explore the gene transcriptional relationships at a much higher resolution, further expanding their knowledge on genetic mechanisms and novel therapeutic targets. Many network construction methods (Bansal et al., 2007; Chai et al., 2014; Karlebach and Shamir, 2008; Marbach et al., 2012; De Smet and Marchal, 2010) have been developed for bulk RNA-seq data, including the popular GGMs. However, they cannot be readily applied to scRNA-seq data (Chen and Mar, 2018; Blencowe et al., 2019). Though both are count data, scRNA-seq data are inherently more sparse with many inflated zero counts. Hence, traditional data processing and normalization methods (e.g., log-transformation) that work well for bulk RNA-seq data may no longer work for the scRNA-seq data (Townes et al., 2019). The excessive amount of zeros in the scRNA-seq data are due to either technical artifacts (false zeros), known as ‘‘dropouts” (Hicks et al., 2018), or biological absence of expression (true zeros) (Jiang et al., 2017; Dong and Jiang, 2019). Only the technical artifacts need to be imputed in the downstream analysis. However, there is no experimental way to differentiate the two types of zeros which challenges the pre-process and analysis of scRNA-seq data.

Furthermore, most existing GRN models for scRNA-seq data are developed for homogeneous cell population under one condition (Matsumoto et al., 2017; Chan et al., 2017; Aibar et al., 2017; Woodhouse et al., 2018; Papili Gao et al., 2018; Sanchez-Castillo et al., 2018; Iacono et al., 2018; Chiu et al., 2018; Deshpande et al., 2019; Aubin-Frankowski and Vert, 2020; Blencowe et al., 2019; Pratapa et al., 2020; Cha and Lee, 2020) with joint GRN methods across multiple conditions just beginning to emerge (Wu et al., 2020). Although the Gaussian copula graph model of Wu et al. (2020) is on joint GRN construction for scRNA-seq data, its major focus is on the novel penalty function that it employs. Additionally, and more importantly, the model directly applies the nonparanormal transformation (Liu et al., 2009) to the count data, which, as mentioned above, may not be appropriate for the single-cell setting (Yoon et al., 2019). To facilitate GGMs and properly employ the Gaussian copula model, for bulk RNA-seq data, Jia et al. (2017) propose to impute and continuize the read counts by learning the posterior distribution under a Bayesian Poisson model framework. Inspired by Jia et al. (2017), we propose to impute and continuize the scRNA-seq count data by a Bayesian zero-inflated Poisson (ZIP) model in which the zero-inflation part accommodates and predicts the dropout events. The choice of the Poisson distribution is supported by the increasing evidence of empirical studies (Townes et al., 2019; Svensson, 2020; Kim et al., 2020). After the dropout zeros are predicted and initially imputed with the Bayesian ZIP model, we further improve the imputation with McImpute (Mongia et al., 2019) to borrow information across genes in realizing that the gene-specific ZIP model does not utilize information across genes. McImpute imputes the dropout events based on low-rank matrix completion, yet it may over-simplify the imputation procedure as it does not employ the gene expression distribution information.

Here, we introduce a comprehensive tool for constructing <u>J</u>oint <u>G</u>ene <u>N</u>etworks with <u>sc</u>RNA-seq (JGNsc) data from different but related conditions. The framework consists of three major steps as shown in Figure 1: (1) a hybrid iterative imputation and continuization procedure; (2) a nonparanormal transformation of the processed data from step (1); and (3) joint GRNs construction which employs the fused lasso technique (Danaher et al., 2014). There are two tuning parameters associated with the model in step (3). Hence, we also investigate the performances of several tuning parameter selection criteria on them, including Akaike information criterion (AIC) (Akaike, 1998), Bayes information criterion (BIC) (Schwarz et al., 1978), Extended Bayesian Information criterion (EBIC) (Chen and Chen, 2008), and the stability approach to regularization selection (StARS) (Liu et al., 2010).

**Figure 1.**
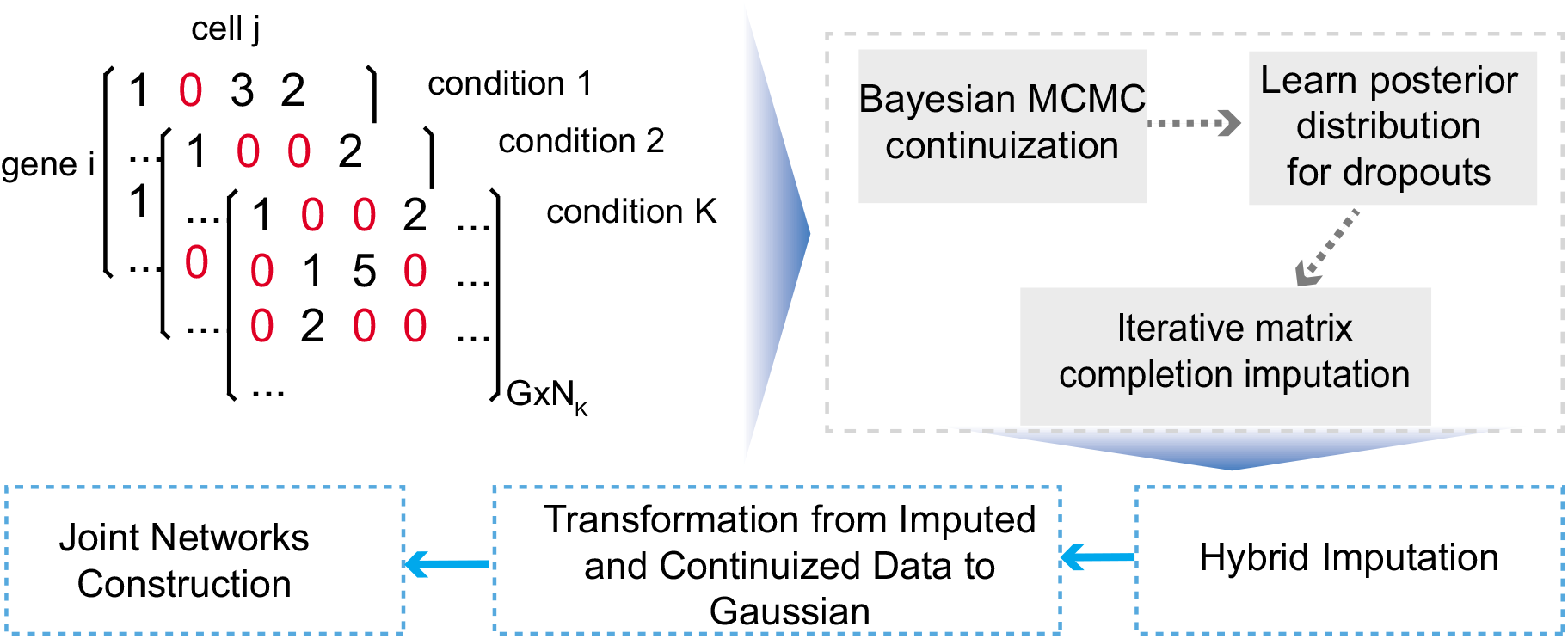
Overview of the JGNsc framework. The JGNsc_Hybrid framework consist of three steps: i) iterative continuization procedure; ii) nonparanormal transformation of the processed data from step i); iii) joint Gaussian graphic gene network construction which employs fused lasso technique and involves several tuning parameters.

This research is motivated by a medulloblastoma scRNA-seq dataset from Hovestadt et al. (2019) and a glioblastoma dataset from Neftel et al. (2019), where single-cell samples from multiple tumor subtypes are available. The research interest is to learn the transcriptional relationships among genes across different tumor subtypes, and thus to guide hypothesis generation for clinical/biological experiments. The rest of the paper is organized as follows: we present the proposed method in Section 2, show results from empirical evaluations and real data analysis in section 3, and conclude in section 4.

## 2. Methods

### 2.1 Hybrid Imputation Procedure for scRNA-seq Data

The proposed Bayesian framework largely follows Jia et al. (2017), in which we replace the Poisson model with the ZIP model to take account the dropout events in scRNA-seq data. For gene *i* (*i* = 1, 2,…, *G*) and cell *j* (*j* = 1, 2,…, *n*), where *G* is the total number of genes and *n* is the total number of cells, let the observed expression read count be *y_ij_*, and the latent variable for indicating the non-dropout event be:

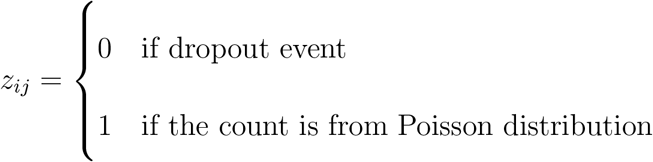

The library size of cell *j* is 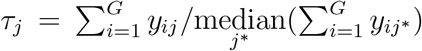. Assuming the gene-specific dropout rate is constant across samples within a homogeneous group, or 1 – *p_i_*, the ZIP model is

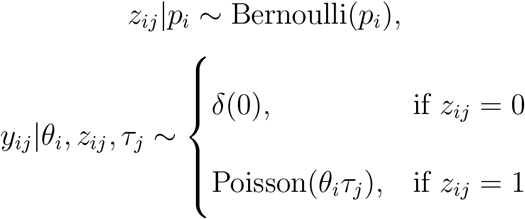

where *δ*(0) is the Dirac delta function with a unit mass concentrated at zero; *θ_i_* is the mean expression factor for gene *i* after being scaled for the library sizes. The priors that we place on *θ_i_* and *p_i_* are

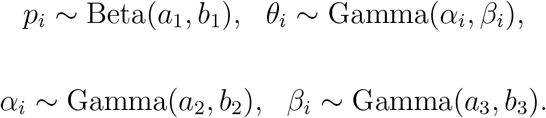

That is, the prior of the mean gene expression *θ_i_* follows a Gamma distribution with two gene-specific parameters *α_i_* and *β_i_*. The priors for *α_i_* and *β_i_* are further assumed to be Gamma distribution, and are a priori independent with hyperparameters *a*_1_, *b*_1_, *a*_2_, *b*_2_, *a*_3_, *b*_3_. The full conditional posterior distributions for the parameters of interest are given below (see supporting information for more details):

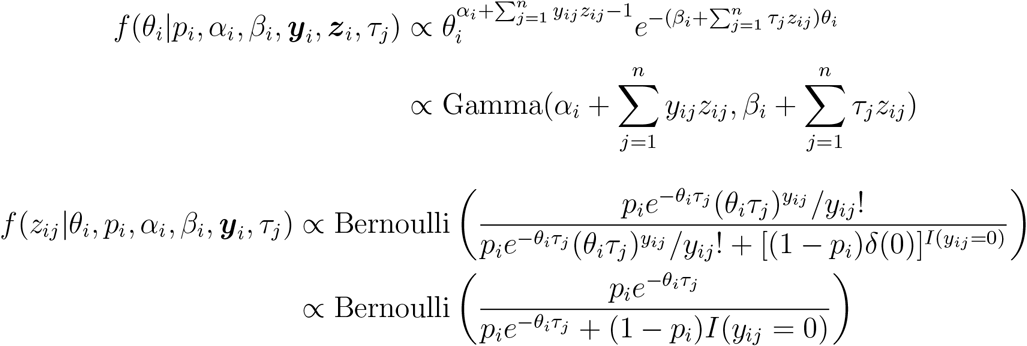

where ***y**_i_* = (*y*_*i*1_, …, *y_in_*) and ***z**_i_* = (*z*_*i*1_, …, *z_in_*). More model details are available in the supporting document. To estimate the posterior distributions, we employ the Metropolis-Hastings within Gibbs sampler MCMC algorithm as follows. Let *t* =1, …,*T* index the iterations in the MCMC procedure. Following the Lemma 1 and Lemma 2 from Jia et al. (2017), we set *a*_2_, *a*_3_ to small positive values, and *b*_2_, *b*_3_ to large numbers, and increase the values of *b*_2_, *b*_3_ in iteration *t* by:

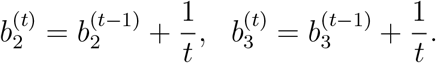

We consider *y_ij_* as a dropout event if the averaged value 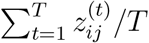 is greater than a pre-chosen threshold, such as 0.75, which we use in the simulated and real data analysis, and impute it to 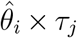, where 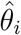 can be the posterior mean or mode of *θ_i_*. We refer the imputed values here as MCMC-imputed values, and let the imputed matrix be 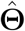. Instead of imputing all zero events as done in Jia et al. (2017) for bulk RNA-seq data, we choose to only impute the zeros that are predicted to be dropouts.

In realizing that 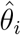 is estimated one gene at a time, we introduce a hybrid imputation procedure that integrates the Bayesian ZIP model and McImpute (Mongia et al., 2019) to further improve the imputation accuracy. McImpute is a low-rank matrix completion method for imputing scRNA-seq data. It borrows expression information from other genes and cells while performing imputation. McImpute has the following objective function:

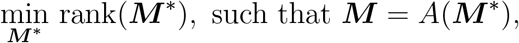

where *A*(·) is a binary mask function indicating whether the element in matrix ***M*** is zero or not with ***M***\* being the complete expression matrix. As suggested by Mongia et al. (2019), this problem can be further converted into a nuclear norm minimization problem and be solved by invoking the Majorization Minimization (Sun et al., 2016). We re-impute the dropout values in 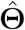 iteratively by Mclmpute (Mongia et al., 2019) as described in Algorithm 1. Within each iteration, *α*% of MCMC-imputed values for dropouts are masked back to zeros. The impact of *α* is investigated in the simulation study.

#### Algorithm 1 JGNsc Hybrid imputation algorithm

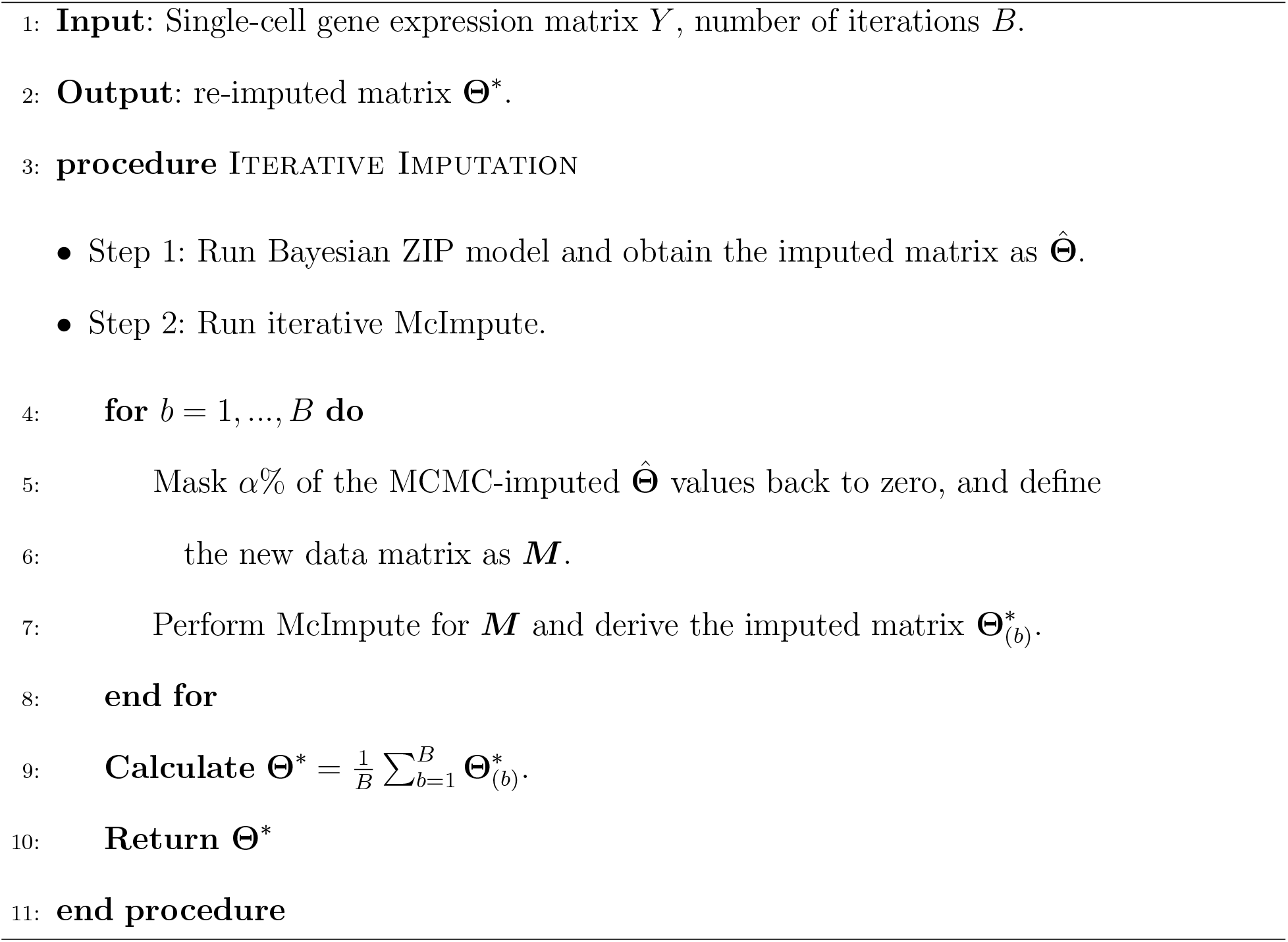

### 2.2 Joint Graphical Lasso Model

Prior to building the joint GGMs, we apply the nonparanormal transformation (Liu et al., 2009) to the final imputed data. The nonparanormal transformation is a popular procedure for pre-processing non-Gaussian distributed data that follow a nonparanormal distribution (Jia et al., 2017; Yoon et al., 2019; Wu et al., 2020). A random vector *X* = (*X*_1_,…, *X_G_*)^*T*^ follows a nonparanormal distribution, or *X* ~ *NPN*(*μ*,Σ, *f*) if there exist functions 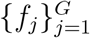 such that *Z* ≡ *f*(*X*) ~ *N*(*μ*, Σ), where *f*(*X*) = (*f*_1_(*X*_1_),…, *f*_*G*_(*X_G_*)). When the *f_j_*’s are monotonic and differentiable, the nonparanormal distribution *NPN*(*μ*, Σ, *f*) is a Gaussian copula, and the conditional independence structure of the original graph is preserved in the precision matrix Ω = Σ^−1^.

After the imputation and the Gaussian transformation, we then employ the Joint Graphical Lasso (JGL) (Danaher et al., 2014), which extends GGMs to simultaneously estimate multiple related precision matrices. For *K* groups of Gaussian data, denote the precision matrix under condition *k* (*k* = 1, …, *K*) as Ω^(*k*)^, with 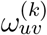 being the (*u, v*)^th^ entry. The task of JGL is to:

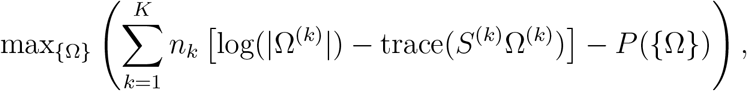

where *n_k_* is the number of observations under condition *k*, *S*^(*k*)^ = (*Y*^(*k*)^)^*T*^*Y*^(*k*)^/*n*_*k*_ is the empirical covariance matrix for the *k*^th^ expression dataset, and *P*({Ω}) is a convex penalty function which is defined as

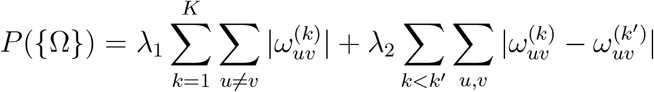

under the fused lasso framework. The penalty function involves two tuning parameters λ_1_ and λ_2_, where λ_1_ ensures a sparsity solution of the graphical lasso model, while λ_2_ enforces the parameters to be shared across conditions.

For tuning parameter selection, many different criteria are available, and the choice of a criterion is largely driven by data and research goals (Wysocki and Rhemtulla, 2019) in practice. There is no one gold standard that has been established for scRNA-seq data. Via simulations, we evaluate four popular tuning parameter selection criteria that are adapted to the JGL models, including AIC, BIC, EBIC, and StARS with detailed definitions given below:

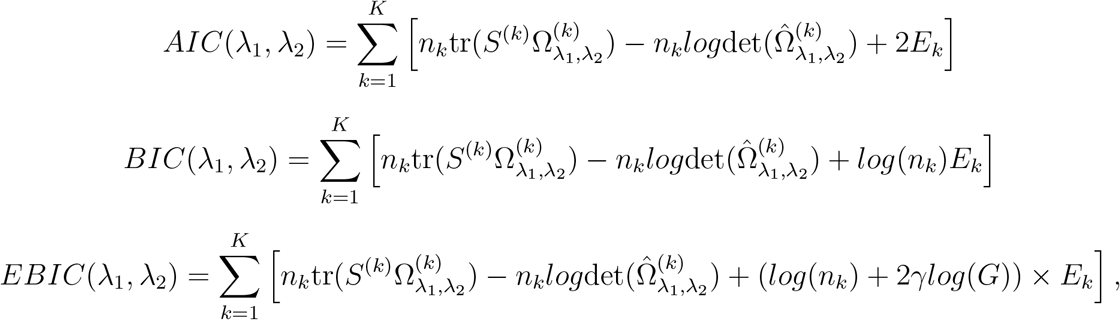

where *E_k_* is the number of non-zero elements in 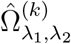, 0 ⩽ *γ* ⩽ 1 is a parameter for EBIC and is set to 0.5 in simulation to control false discovery rate while maintaining positive selection rate. We adapt the StARS method to the JGL models by optimizing one parameter at a time iteratively with details outlined in the supplements.

### 2.3 Simulation of scRNA-seq Data under Different Conditions

To simulate scRNA-seq data under different but related conditions, we first simulate the conditional independence structures based on an inverse nonparanormal transformation algorithm developed by Yahav and Shmueli (2012), which is later applied by Jia et al. (2017) according to the simulation scheme below. For each condition *k* (*k* = 1, 2, …, *K*),

a. Randomly sample from a multivariate Gaussian distribution with a known precision matrix **Ω**^(*k*)^ (Details available in Web supporting information). Denote the random samples by ***X***_1_,…, ***X***_*G*_, where each variable ***X**_i_* = (*X*_*i*1_,…, *X_in_*)^*T*^ consists of *n* realizations.
b. For each variable ***X**_i_*, derive the empirical CDF based on the *n* realizations, and then calculate the cumulative probability for each *X_ij_*.
c. Generate the *k*th scRNA-seq count variables with pre-specified parameters 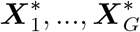 where 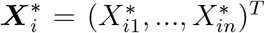 consists of *n* realizations. 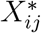 is sampled from a Poisson distribution with true mean level *θ_i_*, which is then assumed to be generated from Gamma distribution. Derive the empirical CDF for each 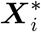, and then calculate the cumulative probability for each 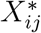.
d. Map the quantiles of the data points in 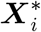, which is generated in (c) to the cumulative probability values calculated from (b). Denote the mapped counts for variable *i* as *Y_ij_*.
e. Based on the zero-inflation parameter distribution, we simulate the random dropout event for each data point *z_ij_*. The final count matrix will be presented as (*z_ij_* × *Y_ij_*).

## 3. Results

This session will show analysis results from our simulation study as well as two real data examples.

### 3.1 Simulation Settings and Results

We simulate scRNA-seq data under *K* = 2 conditions following the strategy described in section 2.3. The number of genes is set to *G* = 100. We vary the sample size, the precision matrix structure, and the dropout rate. Since *n* < *G* and *n* ⩾ *G* are both possible in real single-cell studies, we set *n*_1_ = *n*_2_ = *n* ∈ {50, 100, 200, 300,500}. To evaluate the impact of the mask rate α on the performance of JGNsc, we also vary α at {5%, 10%,…, 95%}. Web Figure 1 shows that JGNsc performs the best when α falls between 10%~ 20%. The reported analysis results from here on are based on the analysis with α of 15%. We investigate the offset of the precision matrix structures in the following three scenarios: i) identical precision matrix structure with different weights; ii) partially identical precision matrix structures and weights; and iii) different precision matrix structures and weights. Here, the weights refer the partial correlations between gene pairs, and the structures refer to the connections between gene pairs. In greater details, for scenario i), all genes are connected with the same structures under two conditions with varying degrees of partial correlations. For scenario ii), we simulate two cases where the first 20 (case 1) or 50 (case 2) genes under the two conditions are set to have different weights and precision matrix structures, but the rest of genes have identical structure and partial correlation values under the two conditions. Each of the above scenarios is simulated for 100 times. To mimic the real scRNA-seq data, we fit the ZIP model to the real single-cell dataset (Hovestadt et al., 2019) using the R package GAMLSS (Rigby and Stasinopoulos, 2005), and the result is shown in Web Figure 2. The estimated parameters from the model are applied to the pre-specified parameters used in steps a)-d) for the simulation study. we simulate *θ_i_* from Gamma(1.5, 0.1); and the non-dropout event probability *p_i_* from Beta(3, 1). Next, we benchmark both the data processing methods and the tuning parameter selection criteria. Benchmark metrics employed in this simulation study include: Pearson correlation between estimated and the true precision matrices, and also sum of squared errors (SSE) of the difference between the two matrices. In addition, the area under the receiver operating characteristic (AUROC) curve and the area under the precision recall (AUPRC) curve are calculated and compared. The data processing methods include: i) NoDropout: No dropout events - we consider this as the best performance that can be achieved; ii) Hybrid: read counts are continuized and imputed by the Bayesian ZIP model and by integrating iterative McImpute for the dropout events; iii) Bayesian: read counts are continuized and imputed by the Bayesian ZIP model only; iv) McImpute: Impute the dropouts by McImpute (Mongia et al., 2019); and v) Observed: the observed raw counts without imputation.

For joint graphical models, tuning parameter selection criteria include AIC, BIC, EBIC and StARS. The results show that the impact of λ_1_, which controls the sparsity of the network, is larger than the impact of *λ*_2_, and the performance of JGNsc results are stable within a reasonable range of *λ*_2_ values. Web Figure 3-7 show that AIC prioritizes the other three criteria, and is used for the real data analysis. Overall, our simulation study shows that the proposed Bayesian continuization and imputation procedure is powerful for improving the downstream gene network construction (Figure 2, Web Figure 8-10).

**Figure 2.**
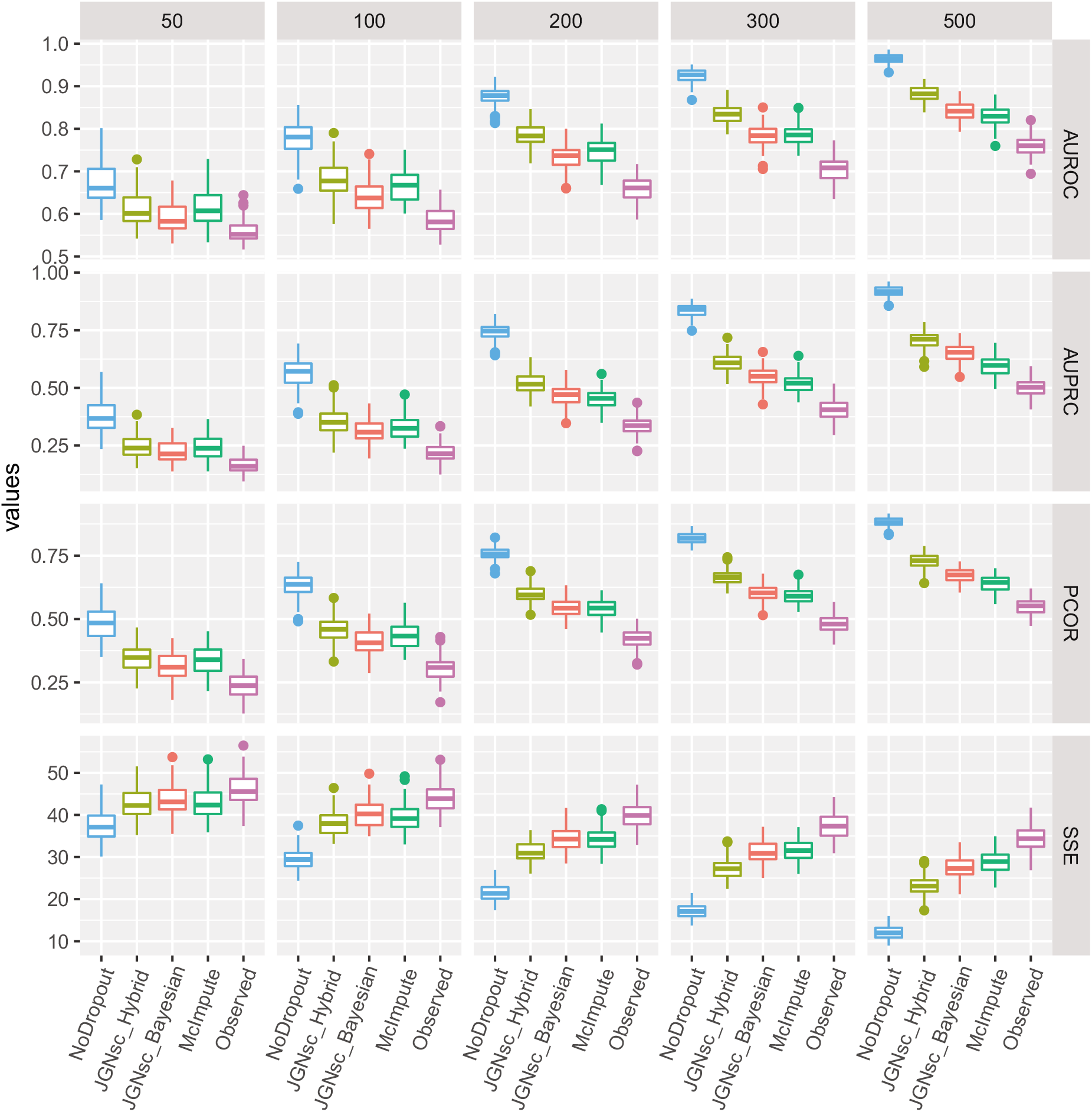
Benchmark of data processing methods with sample size varying. As the sample size increases, the overall performance is improved for each method. The JGNsc_Hybrid method outperforms other benchmark methods in terms of all four metrics for the joint network partial correlation. The performance of JGNsc_Bayesian (without iteration) becomes better and excel McImpute when the sample size is large (500).

### 3.2 Analysis of scRNA-seq data from medulloblastoma and glioblastoma (GBM)

Medulloblastoma (MB) is the most common pediatric brain tumor for which oncogenic drivers, including specific transcriptional factors (TFs), have been well defined. Oncogenes essential in MB tumorigenesis include Myc, a key oncogenic TF for Group 3 MBs (Northcott et al., 2011), and Otx2, a TF functionally interacting with Myc in Group 3 MBs and playing essential oncogenic roles in Group 4 MBs (Bunt et al., 2011; Lu et al., 2017; Boulay et al., 2017).

As a key oncoprotein in a variety of human cancers, Myc is believed to play essential roles in controlling the transcription of a large number genes and regulating multiple cellular processes in cancer cells, including, most prominently, protein biogenesis and cell metabolism (Van Riggelen et al., 2010; Dang, 2012). We postulate that JGNsc can help delineate the roles of Myc and Otx2 in regulating gene transcription as well as their functional interaction in different subtypes of MB cells. Utilizing the MB scRNA-seq data set (GSE119926) by Hovestadt et al. (2019), we selected samples from a subset of 17 individuals that were grouped into two subtypes of medulloblastoma based on their molecular profiles (Hovestadt et al., 2019): Group 3 or Group 4, or another subset of cells that falls between these two subtypes (Intermediate group) (Web Figure 11).

We continuize and impute the dropout events in the scRNA-seq data using a selected set of ~ 6K genes (genes of interest and genes with non-zero counts in at least 300 cells in each group), and use the enzyme-related genes from mammalian metabolic enzyme database (Corcoran et al., 2017) for the joint network inference. AIC is used to select the tuning parameter values for the JGNsc model. We visualize the Myc-connected and Otx2-connected genes under different conditions in Figure 3.

**Figure 3.**
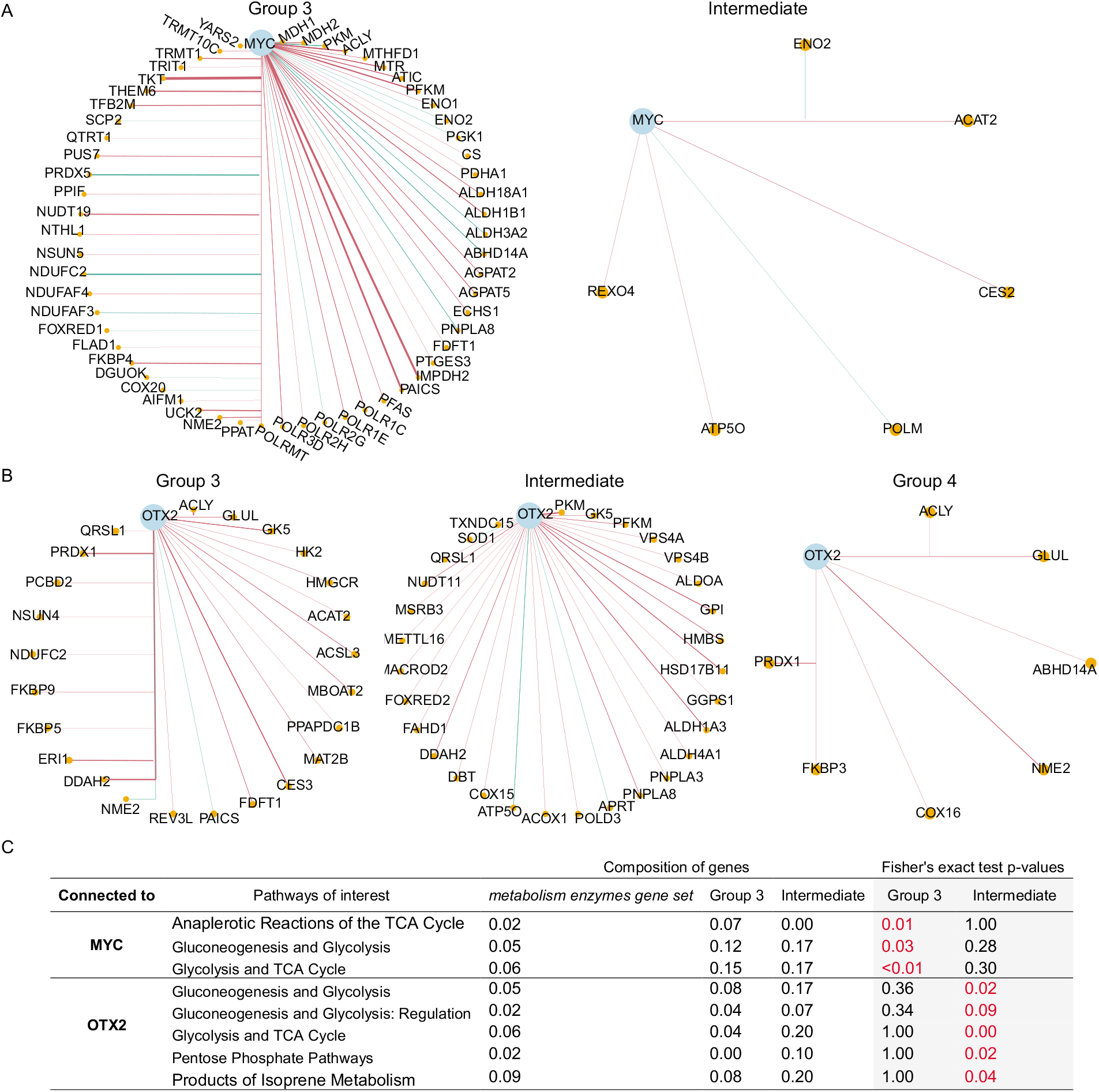
JGNsc networks for Medulloblastoma data. In a network, each node is a gene and each edge shows the partial correlation between a pair of genes. If the partial correlation is zero, there is no connection between this pair of genes. Further, if two genes are connected by a red line, then their partial correlation is positive; otherwise if they are connected by a green line, their partial correlation is negative. (A) Network visualization of JGNsc results for Myc related genes from purine enzymes metabolism pathway. Connections between non-MYC related genes are not shown. (B) Network visualization of JGNsc results for Otx2 related genes from purine enzymes metabolism pathway. Connections between non-Otx2 related genes are not shown. (C) GSEA for metabolism enzymes genes connected to Myc or Otx2. Fisher’s exact test was performed for each group and each pathway respectively. The pathways with significant p-value for Fisher’s exact test for either Myc or Otx2 connected gene set under a condition. The complete results are available at supporting information Web Figure 12.

For Myc-connected genes (Figure 3A), the network for group 3 is denser compared to the intermediate group, while no connection is detected for Myc in the group 4 samples, in agreement with the prominent roles of Myc in Group 3 MBs (Northcott et al., 2011; Hovestadt et al., 2019). Interestingly, the parallel analysis on Otx2 leads to the following two observations: (1) Otx2 is connected to metabolic genes in the intermediate group of MB cells but less so in the Group 4 MB cells; and (2) in Group 3 MBs, where Otx2 is thought to be functionally cooperating with Myc (Lu et al., 2017; Bunt et al., 2011; Boulay et al., 2017), its connection to metabolic genes was distinct from Myc’s, highlighting the unique link between Myc and metabolism in Group 3 MB cells (Figure 3B). Further Gene Set Enrichment Analysis (GSEA) based on the previously defined metabolic pathway (Corcoran et al., 2017) reveals that overlapping yet distinct metabolic pathways were connected to each of the two TFs (Figure 3C). Collectively, these results from the JGNsc analysis suggest that the role of Myc in regulating the expression of metabolic genes is MB-subtype-dependent, and that Myc and Otx2 likely play diverse roles in regulating the transcription of metabolic genes.

Next, we extend the JGNsc analysis to a scRNA-seq data from another type of brain tumor, GBM. With a median survival of about 12-15 months, GBM is the most common and lethal brain tumor, due to its intratumoral heterogeneity and treatment resistance. It is believed that these features of GBMs are largely due to the existence of a subset of tumor initiating cells - GBM stem cells (GSCs) (Lathia et al., 2015), and that Myc, a TF in regulating the expression of metabolic genes, is essential for sustaining this population of tumor cells (Wang et al., 2008, 2017).

To test the connection between Myc and the expression of metabolic enzyme-encoding genes in GBM tumor cells exhibiting prevalent heterogeneity, we apply JGNsc to a subset of GBM scRNA-seq data (GSE131928) from Neftel et al. (2019). In agreement with the heterogeneous nature of GBM tumor cells, GBM scRNA-seq samples are classified into four subgroups with distinct gene expression profiles: neural-progenitor-like (NPC-like), oligodendrocyte-progenitor-like (OPC-like), astrocyte-like (AC-like), and mesenchymal-like (MES-like) tumor cells (Neftel et al., 2019). For simplicity, we only take the malignant single-cell samples from adult GBM patients sequenced by the Smart-seq2 protocol (Picelli et al., 2013), and classify these tumor cells into 4 subgroups as defined by the original study (Web Figure 13). This results in 1213 OPC-like cells, 1267 NPC-like cells, 1262 AC-like cells, and 637 MES-like cells. Our analysis finds that there are over 20% of cells expressing Myc in each of the subgroup of GBM cells (Web Figure 13B, D) in agreement with the essential roles of Myc in maintaining a subset of GSCs (Wang et al., 2008, 2017). Further joint network analysis on Myc’s connection to metabolism enzyme-encoding genes reveals distinct connection profiles in the four subgroups of GBM cells. MES-like tumor cells demonstrate the most connected gene, followed by AC-like and OPC-like cells, while only one gene is found to be connected to Myc in the OPC-like subgroup (Web Figure 14). Subsequent GSEA identifies tumor cell subgroup-dependent metabolic pathway’s connection to Myc: *Anaplerotic Reactions of the TCA Cycle*, *Folic Acid Metabolism* are enriched in the MES-like group, *Lipid Metabolism* is up-regulated in the OPC-like group, while *Pyrimidine Metabolism* is enriched in the AC-like group (Web Figure 15). Echoing the findings from the MB cells, these results suggest that Myc’s roles in regulating metabolic processes and the mechanisms underlying Myc-mediated GSC maintenance likely vary in different subgroups of GBM cells.

## 4. Discussion

We propose an integrated framework JGNsc to construct joint gene networks using scRNA-seq data under different conditions. The JGNsc framework consists of three major steps: first, continuize and impute scRNA-seq data using a MCMC procedure; second, transform the data into Gaussian form using nonparanormal distribution; and third, jointly construct gene network using JGL. The novelty of our paper lies in the MCMC data continuization step for scRNA-seq data, where iterative matrix completion imputation is implemented. This step helps to correct the posterior distribution of non-dropout event, and in turn improve the expression estimation of the dropouts.

In our simulation, we also demonstrate that a proper imputation of scRNA-seq data would greatly improve the downstream joint network construction performance. In addition, JGNsc performs consistently better than the other benchmark methods when we vary the sample size and data structure. By transforming the estimated precision matrices into the partial correlation matrices, our method also allows the investigation of the relative magnitude and direction of gene-wise connections.

As future research, we may adopt the following penalty function:

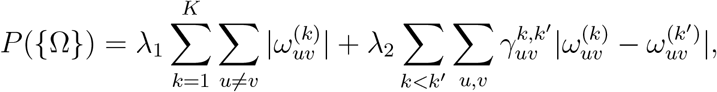

where 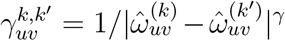 is the weight assigned for the pair of 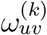 and 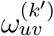 to constrain the level of their penalty, and γ is a positive constant for adjustment of the adaptive weight matrix. The values 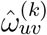 and 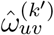 are the estimated precision matrices elements from *K* naive GGMs. The idea is motivated by the adaptive lasso of Zou (2006) which has the oracle properties and allows consistent variable selection, and the condition-adaptive fused graphical lasso for bulk RNA-seq data of Lyu et al. (2018). By imposing a gene-pair-specific binary weight between conditions, Lyu et al. (2018) only put constrains on edges that are tested to be non-differentially co-expressed. Such strategies could potentially further improve the network constructions.

## Acknowledgements

The authors thank for the suggestions from reviewers.

## Supplementary Materials

Supplemental figures and tables are available with this paper at the Biometrics website on Wiley Online Library.

## Code and Data Availability

JGNsc framework R code is available online at github: https://github.com/meichendong/JGNsc. The raw data from Hovestadt et al. (2019) is available at Gene Expression Omnibus: GSE119926. The raw data from Neftel et al. (2019) is available at Gene Expression Omnibus: GSE131928.

## References

Aibar, S., González-Blas, C. B., Moerman, T., Imrichova, H., Hulselmans, G., Rambow, F., Marine, J.-C., Geurts, P., Aerts, J., van den Oord, J., et al. (2017). Scenic: single-cell regulatory network inference and clustering. Nature methods 14, 1083–1086.

Akaike, H. (1998). Information theory and an extension of the maximum likelihood principle. In Selected papers of hirotugu akaike, pages 199–213. Springer.

Allocco, D. J., Kohane, I. S., and Butte, A. J. (2004). Quantifying the relationship between co-expression, co-regulation and gene function. BMC bioinformatics 5, 1–10.

Aubin-Frankowski, P.-C. and Vert, J.-P. (2020). Gene regulation inference from single-cell rna-seq data with linear differential equations and velocity inference. Bioinformatics 36, 4774–4780.

Ballouz, S., Verleyen, W., and Gillis, J. (2015). Guidance for rna-seq co-expression network construction and analysis: safety in numbers. Bioinformatics 31, 2123–2130.

Bansal, M., Belcastro, V., Ambesi-Impiombato, A., and Di Bernardo, D. (2007). How to infer gene networks from expression profiles. Molecular systems biology 3, 78.

Blencowe, M., Arneson, D., Ding, J., Chen, Y.-W., Saleem, Z., and Yang, X. (2019). Network modeling of single-cell omics data: challenges, opportunities, and progresses. Emerging topics in life sciences 3, 379–398.

Boulay, G., Awad, M. E., Riggi, N., Archer, T. C., Iyer, S., Boonseng, W. E., Rossetti, N. E., Naigles, B., Rengarajan, S., Volorio, A., et al. (2017). Otx2 activity at distal regulatory elements shapes the chromatin landscape of group 3 medulloblastoma. Cancer discovery 7, 288–301.

Buil, A., Brown, A. A., Lappalainen, T., Viñ uela, A., Davies, M. N., Zheng, H.-F., Richards, J. B., Glass, D., Small, K. S., Durbin, R., et al. (2015). Gene-gene and gene-environment interactions detected by transcriptome sequence analysis in twins. Nature genetics 47, 88–91.

Bunt, J., Hasselt, N. E., Zwijnenburg, D. A., Koster, J., Versteeg, R., and Kool, M. (2011). Joint binding of otx2 and myc in promotor regions is associated with high gene expression in medulloblastoma. PloS one 6, e26058.

Cai, T. T., Li, H., Liu, W., and Xie, J. (2016). Joint estimation of multiple high-dimensional precision matrices. Statistica Sinica 26, 445.

Cha, J. and Lee, I. (2020). Single-cell network biology for resolving cellular heterogeneity in human diseases. Experimental & Molecular Medicine pages 1–11.

Chai, L. E., Loh, S. K., Low, S. T., Mohamad, M. S., Deris, S., and Zakaria, Z. (2014). A review on the computational approaches for gene regulatory network construction. Computers in biology and medicine 48, 55–65.

Chan, T. E., Stumpf, M. P., and Babtie, A. C. (2017). Gene regulatory network inference from single-cell data using multivariate information measures. Cell systems 5, 251–267.

Chen, J. and Chen, Z. (2008). Extended bayesian information criteria for model selection with large model spaces. Biometrika 95, 759–771.

Chen, S. and Mar, J. C. (2018). Evaluating methods of inferring gene regulatory networks highlights their lack of performance for single cell gene expression data. BMC bioinformatics 19, 1–21.

Chiu, Y.-C., Hsiao, T.-H., Wang, L.-J., Chen, Y., and Shao, Y.-H. J. (2018). scdnet: a computational tool for single-cell differential network analysis. BMC systems biology 12, 124.

Chun, H., Zhang, X., and Zhao, H. (2015). Gene regulation network inference with joint sparse gaussian graphical models. Journal of Computational and Graphical Statistics 24, 954–974.

Corcoran, C. C., Grady, C. R., Pisitkun, T., Parulekar, J., and Knepper, M. A. (2017). From 20th century metabolic wall charts to 21st century systems biology: database of mammalian metabolic enzymes. American Journal of Physiology-Renal Physiology 312, F533–F542.

Danaher, P., Wang, P., and Witten, D. M. (2014). The joint graphical lasso for inverse covariance estimation across multiple classes. Journal of the Royal Statistical Society: Series B (Statistical Methodology) 76, 373–397.

Dang, C. V. (2012). Myc on the path to cancer. Cell 149, 22–35.

De Smet, R. and Marchal, K. (2010). Advantages and limitations of current network inference methods. Nature Reviews Microbiology 8, 717–729.

Deshpande, A., Chu, L.-F., Stewart, R., and Gitter, A. (2019). Network inference with granger causality ensembles on single-cell transcriptomic data. BioRxiv page 534834.

Dong, M. and Jiang, Y. (2019). Single-cell allele-specific gene expression analysis. In Computational Methods for Single-Cell Data Analysis, pages 155–174. Springer.

Faith, J. J., Hayete, B., Thaden, J. T., Mogno, I., Wierzbowski, J., Cottarel, G., Kasif, S., Collins, J. J., and Gardner, T. S. (2007). Large-scale mapping and validation of escherichia coli transcriptional regulation from a compendium of expression profiles. PLoS biol 5, e8.

Geng, S., Yan, M., Kolar, M., and Koyejo, O. (2019). Partially linear additive gaussian graphical models. arXiv preprint arXiv:1906.03362 .

Grimes, T., Potter, S. S., and Datta, S. (2019). Integrating gene regulatory pathways into differential network analysis of gene expression data. Scientific reports 9, 1–12.

Ha, M. J., Baladandayuthapani, V., and Do, K.-A. (2015). Dingo: differential network analysis in genomics. Bioinformatics 31, 3413–3420.

Hallac, D., Park, Y., Boyd, S., and Leskovec, J. (2017). Network inference via the time-varying graphical lasso. In Proceedings of the 23rd ACM SIGKDD International Conference on Knowledge Discovery and Data Mining, pages 205–213.

Haslbeck, J. and Waldorp, L. J. (2015). mgm: Estimating time-varying mixed graphical models in high-dimensional data. arXiv preprint arXiv:1510.06871.

Hicks, S. C., Townes, F. W., Teng, M., and Irizarry, R. A. (2018). Missing data and technical variability in single-cell rna-sequencing experiments. Biostatistics 19, 562–578.

Hovestadt, V., Smith, K. S., Bihannic, L., Filbin, M. G., Shaw, M. L., Baumgartner, A., DeWitt, J. C., Groves, A., Mayr, L., Weisman, H. R., et al. (2019). Resolving medulloblastoma cellular architecture by single-cell genomics. Nature 572, 74–79.

Huang, F., Chen, S., and Huang, S.-J. (2017). Joint estimation of multiple conditional gaussian graphical models. IEEE transactions on neural networks and learning systems 29, 3034–3046.

Iacono, G., Mereu, E., Guillaumet-Adkins, A., Corominas, R., Cuscó, I., Rodríguez-Esteban, G., Gut, M., Pérez-Jurado, L. A., Gut, I., and Heyn, H. (2018). bigscale: an analytical framework for big-scale single-cell data. Genome research 28, 878–890.

Jia, B. and Liang, F. (2019). Fast hybrid bayesian integrative learning of multiple gene regulatory networks for type 1 diabetes. Biostatistics.

Jia, B., Xu, S., Xiao, G., Lamba, V., and Liang, F. (2017). Learning gene regulatory networks from next generation sequencing data. Biometrics 73, 1221–1230.

Jiang, Y., Zhang, N. R., and Li, M. (2017). Scale: modeling allele-specific gene expression by single-cell rna sequencing. Genome biology 18, 74.

Karlebach, G. and Shamir, R. (2008). Modelling and analysis of gene regulatory networks. Nature Reviews Molecular Cell Biology 9, 770–780.

Kim, T., Zhou, X., and Chen, M. (2020). Demystifying” drop-outs” in single cell umi data. bioRxiv.

Langfelder, P. and Horvath, S. (2008). Wgcna: an r package for weighted correlation network analysis. BMC bioinformatics 9, 559.

Lathia, J. D., Mack, S. C., Mulkearns-Hubert, E. E., Valentim, C. L., and Rich, J. N. (2015). Cancer stem cells in glioblastoma. Genes & development 29, 1203–1217.

Lee, S., Liang, F., Cai, L., and Xiao, G. (2018). A two-stage approach of gene network analysis for high-dimensional heterogeneous data. Biostatistics 19, 216–232.

Liu, H., Lafferty, J., and Wasserman, L. (2009). The nonparanormal: Semiparametric estimation of high dimensional undirected graphs. Journal of Machine Learning Research 10,.

Liu, H., Roeder, K., and Wasserman, L. (2010). Stability approach to regularization selection (stars) for high dimensional graphical models. In Advances in neural information processing systems, pages 1432–1440.

Lu, Y., Labak, C. M., Jain, N., Purvis, I. J., Guda, M. R., Bach, S. E., Tsung, A. J., Asuthkar, S., and Velpula, K. K. (2017). Otx2 expression contributes to proliferation and progression in myc-amplified medulloblastoma. American journal of cancer research 7, 647.

Lyu, Y., Xue, L., Zhang, F., Koch, H., Saba, L., Kechris, K., and Li, Q. (2018). Condition-adaptive fused graphical lasso (cfgl): An adaptive procedure for inferring condition-specific gene co-expression network. PLoS computational biology 14, e1006436.

Marbach, D., Costello, J. C., Küffner, R., Vega, N. M., Prill, R. J., Camacho, D. M., Allison, K. R., Kellis, M., Collins, J. J., and Stolovitzky, G. (2012). Wisdom of crowds for robust gene network inference. Nature methods 9, 796–804.

Margolin, A. A., Nemenman, I., Basso, K., Wiggins, C., Stolovitzky, G., Dalla Favera, R., and Califano, A. (2006). Aracne: an algorithm for the reconstruction of gene regulatory networks in a mammalian cellular context. In BMC bioinformatics, volume 7, page S7. Springer.

Matsumoto, H., Kiryu, H., Furusawa, C., Ko, M. S., Ko, S. B., Gouda, N., Hayashi, T., and Nikaido, I. (2017). Scode: an efficient regulatory network inference algorithm from single-cell rna-seq during differentiation. Bioinformatics 33, 2314–2321.

Mongia, A., Sengupta, D., and Majumdar, A. (2019). Mcimpute: Matrix completion based imputation for single cell rna-seq data. Frontiers in genetics 10, 9.

Neftel, C., Laffy, J., Filbin, M. G., Hara, T., Shore, M. E., Rahme, G. J., Richman, A. R., Silverbush, D., Shaw, M. L., Hebert, C. M., et al. (2019). An integrative model of cellular states, plasticity, and genetics for glioblastoma. Cell 178, 835–849.

Northcott, P. A., Korshunov, A., Witt, H., Hielscher, T., Eberhart, C. G., Mack, S., Bouffet, E., Clifford, S. C., Hawkins, C. E., French, P., et al. (2011). Medulloblastoma comprises four distinct molecular variants. Journal of clinical oncology 29, 1408.

Oates, C. J., Korkola, J., Gray, J. W., Mukherjee, S., et al. (2014). Joint estimation of multiple related biological networks. The Annals of Applied Statistics 8, 1892–1919.

Papili Gao, N., Ud-Dean, S. M., Gandrillon, O., and Gunawan, R. (2018). Sincerities: inferring gene regulatory networks from time-stamped single cell transcriptional expression profiles. Bioinformatics 34, 258–266.

Peng, H., Long, F., and Ding, C. (2005). Feature selection based on mutual information criteria of max-dependency, max-relevance, and min-redundancy. IEEE Transactions on pattern analysis and machine intelligence 27, 1226–1238.

Picelli, S., Bjorklund, Å. K., Faridani, O. R., Sagasser, S., Winberg, G., and Sandberg, R. (2013). Smart-seq2 for sensitive full-length transcriptome profiling in single cells. Nature methods 10, 1096–1098.

Pratapa, A., Jalihal, A. P., Law, J. N., Bharadwaj, A., and Murali, T. (2020). Benchmarking algorithms for gene regulatory network inference from single-cell transcriptomic data. Nature Methods 17, 147–154.

Qiu, H., Han, F., Liu, H., and Caffo, B. (2016). Joint estimation of multiple graphical models from high dimensional time series. Journal of the Royal Statistical Society. Series B, Statistical Methodology 78, 487.

Ramirez, A. K., Dankel, S. N., Rastegarpanah, B., Cai, W., Xue, R., Crovella, M., Tseng, Y.-H., Kahn, C. R., and Kasif, S. (2020). Single-cell transcriptional networks in differentiating preadipocytes suggest drivers associated with tissue heterogeneity. Nature Communications 11, 1–9.

Rigby, R. A. and Stasinopoulos, D. M. (2005). Generalized additive models for location, scale and shape,(with discussion). Applied Statistics 54, 507–554.

Sales, G. and Romualdi, C. (2011). parmigene—a parallel r package for mutual information estimation and gene network reconstruction. Bioinformatics 27, 1876–1877.

Sanchez-Castillo, M., Blanco, D., Tienda-Luna, I. M., Carrion, M., and Huang, Y. (2018). A bayesian framework for the inference of gene regulatory networks from time and pseudo-time series data. Bioinformatics 34, 964–970.

Schwarz, G. et al. (1978). Estimating the dimension of a model. The annals of statistics 6, 461–464.

Sun, Y., Babu, P., and Palomar, D. P. (2016). Majorization-minimization algorithms in signal processing, communications, and machine learning. IEEE Transactions on Signal Processing 65, 794–816.

Svensson, V. (2020). Droplet scrna-seq is not zero-inflated. Nature Biotechnology pages 1–4.

Townes, F. W., Hicks, S. C., Aryee, M. J., and Irizarry, R. A. (2019). Feature selection and dimension reduction for single-cell rna-seq based on a multinomial model. Genome biology 20, 1–16.

Trapnell, C., Cacchiarelli, D., Grimsby, J., Pokharel, P., Li, S., Morse, M., Lennon, N. J., Livak, K. J., Mikkelsen, T. S., and Rinn, J. L. (2014). The dynamics and regulators of cell fate decisions are revealed by pseudotemporal ordering of single cells. Nature biotechnology 32, 381.

van Dam, S., Vosa, U., van der Graaf, A., Franke, L., and de Magalhaes, J. P. (2018). Gene co-expression analysis for functional classification and gene-disease predictions. Briefings in bioinformatics 19, 575–592.

Van de Sande, B., Flerin, C., Davie, K., De Waegeneer, M., Hulselmans, G., Aibar, S., Seurinck, R., Saelens, W., Cannoodt, R., Rouchon, Q., et al. (2020). A scalable scenic workflow for single-cell gene regulatory network analysis. Nature Protocols 15, 2247–2276.

Van Riggelen, J., Yetil, A., and Felsher, D. W. (2010). Myc as a regulator of ribosome biogenesis and protein synthesis. Nature Reviews Cancer 10, 301–309.

Wang, J., Wang, H., Li, Z., Wu, Q., Lathia, J. D., McLendon, R. E., Hjelmeland, A. B., and Rich, J. N. (2008). c-myc is required for maintenance of glioma cancer stem cells. PloS one 3, e3769.

Wang, X., Yang, K., Xie, Q., Wu, Q., Mack, S. C., Shi, Y., Kim, L. J., Prager, B. C., Flavahan, W. A., Liu, X., et al. (2017). Purine synthesis promotes maintenance of brain tumor initiating cells in glioma. Nature neuroscience 20, 661–673.

Wolfe, C. J., Kohane, I. S., and Butte, A. J. (2005). Systematic survey reveals general applicability of” guilt-by-association” within gene coexpression networks. BMC bioinformatics 6, 227.

Woodhouse, S., Piterman, N., Wintersteiger, C. M., Göttgens, B., and Fisher, J. (2018). Scns: a graphical tool for reconstructing executable regulatory networks from single-cell genomic data. BMC systems biology 12, 1–7.

Wu, N., Huang, J., Zhang, X.-F., Ou-Yang, L., He, S., Zhu, Z., and Xie, W. (2019). Weighted fused pathway graphical lasso for joint estimation of multiple gene networks. Frontiers in genetics 10, 623.

Wu, N., Yin, F., Ou-Yang, L., Zhu, Z., and Xie, W. (2020). Joint learning of multiple gene networks from single-cell gene expression data. Computational and Structural Biotechnology Journal 18, 2583–2595.

Wysocki, A. C. and Rhemtulla, M. (2019). On penalty parameter selection for estimating network models. Multivariate behavioral research pages 1–15.

Yahav, I. and Shmueli, G. (2012). On generating multivariate poisson data in management science applications. Applied Stochastic Models in Business and Industry 28, 91–102.

Yang, S., Lu, Z., Shen, X., Wonka, P., and Ye, J. (2015). Fused multiple graphical lasso. SIAM Journal on Optimization 25, 916–943.

Yoon, G., Gaynanova, I., and Muller, C. L. (2019). Microbial networks in spring-semi-parametric rank-based correlation and partial correlation estimation for quantitative microbiome data. Frontiers in genetics 10, 516.

Zhang, A., Cai, B., Hu, W., Jia, B., Liang, F., Wilson, T. W., Stephen, J. M., Calhoun, V. D., and Wang, Y.-P. (2019). Joint bayesian-incorporating estimation of multiple gaussian graphical models to study brain connectivity development in adolescence. IEEE transactions on medical imaging 39, 357–365.

Zhu, Y. and Koyejo, O. (2018). Clustered fused graphical lasso. In UAI, pages 487–496.

Zou, H. (2006). The adaptive lasso and its oracle properties. Journal of the American statistical association 101, 1418–1429.

