## Supplement Materials for "Joint Gene Network Construction by Single-Cell RNA Sequencing Data"

Biometrics xx, 1–8 DOI: pending......
xxxxx 2021

### Supporting Information for JGNsc

**Meichen Dong[1], Yiping He[2], Yuchao Jiang [1,3], Fei Zou[1,3,*]**

[1]Department of Biostatistics, University of North Carolina at Chapel Hill, Chapel Hill, North Carolina, U.S.A.

[2]Department of Pathology, Duke University, Durham, North Carolina, U.S.A.

[3]Department of Genetics, University of North Carolina at Chapel Hill, Chapel Hill, North Carolina, U.S.A.

This paper has been submitted for consideration for publication in *Biometrics*

#### 1. Details of the JGNsc Bayesian framework

For gene $i$, cell $j$, assume the observed gene expression read count is $y_{ij}$, the library size factor of cell $j$ is $\tau_j = \sum_{i=1}^{G} y_{ij}/\underset{j^*}{\text{median}}(\sum_{i=1}^{G} y_{ij^*})$, the non-dropout indicator is:

$$z_{ij} = \begin{cases} 0 & \text{if dropout event} \\ 1 & \text{if the count is from Poisson distribution} \end{cases}$$

, the non-dropout event probability is $p_i$. The model specification is as below:

$$z_{ij}|p_i \sim \text{Bernoulli}(p_i)$$

$$y_{ij}|\theta_i, z_{ij}, \tau_j \sim \begin{cases} \delta(0), & \text{if } z_{ij} = 0 \\ \text{Poisson}(\theta_i \tau_j), & \text{if } z_{ij} = 1 \end{cases}$$

where $\delta(0)$ is the Dirac delta function with a unit mass concentrated at zero. $\theta_i$ is the mean expression factor for gene $i$.

For observed zero counts $y_{ij} = 0$,

$$f(z_{ij}|p_i) = p_i^{z_{ij}}(1-p_i)^{1-z_{ij}}, \quad 0 < p_i < 1.$$

$$f_1(y_{ij}, z_{ij}|\theta_i, p_i, \tau_j) = [(1-p_i)\delta(0)]^{1-z_{ij}} \times p_i^{z_{ij}} \times \left[\frac{e^{-\theta_i\tau_j}(\theta_i\tau_j)^{y_{ij}}}{y_{ij}!}\right]^{z_{ij}}, \quad z_{ij} = 0, 1$$

For observed non-zero counts $y_{ij} > 0$,

$$f_2(y_{ij}|\theta_i, p_i, \tau_j) = p_i \times \frac{e^{-\theta_i\tau_j}(\theta_i\tau_j)^{y_{ij}}}{y_{ij}!}, \quad y_{ij} = 1, 2, ...; \; z_{ij} = 1.$$

Prior distributions of parameters in the model are specified as:

$$p_i \sim \text{Beta}(a_1, b_1), \quad f(p_i|a_1, b_1) = \frac{1}{B(a_1, b_1)} p_i^{a_1-1}(1-p_i)^{b_1-1}, \; 0 < p_i < 1$$

$$\theta_i \sim \text{Gamma}(\alpha_i, \beta_i), \quad f(\theta_i|\alpha_i, \beta_i) = \frac{\beta_i^{\alpha_i}}{\Gamma(\alpha_i)} \theta_i^{\alpha_i-1} exp(-\beta_i\theta_i)$$

$$\alpha_i \sim \text{Gamma}(a_2, b_2), \quad f(\alpha_i|a_2, b_2) = \frac{b_2^{a_2}}{\Gamma(a_2)} \alpha_i^{a_2-1} exp(-b_2\alpha_i), \quad \alpha_i > 0$$

$$\beta_i \sim \text{Gamma}(a_3, b_3), \quad f(\beta_i|a_3, b_3) = \frac{b_3^{a_3}}{\Gamma(a_3)} \beta_i^{a_3-1} exp(-b_3\beta_i), \quad \beta_i > 0$$

, where $a_1, b_1, a_2, b_2, a_3, b_3$ are prior hyperparameters.

For all the $n$ cells under a certain condition, the joint likelihood for gene $i$ can be written as:

$$
\begin{aligned}
&f(\boldsymbol{y}_i, \boldsymbol{z}_i, \theta_i, p_i, \tau_j, \alpha_i, \beta_i) \\
&= \left[ \prod_{y_{ij}=0} f_1(y_{ij}, z_{ij}|\theta_i, p_i, \tau_j) \prod_{y_{ij}>0} f_2(y_{ij}|\theta_i, p_i, \tau_j) \right] \times f(p_i|a_1, b_1) \times f(\theta_i|\alpha_i, \beta_i) \\
&\quad \times f(\alpha_i|a_2, b_2) \times f(\beta_i|a_3, b_3) \\
&= \left[ \prod_{j=1}^{n} \left( [(1-p_i)\delta(0)]^{1-z_{ij}} \times p_i^{z_{ij}} \left[ \frac{e^{-\theta_i\tau_j}(\theta_i\tau_j)^{y_{ij}}}{y_{ij}!} \right]^{z_{ij}} \right)^{I(y_{ij}=0)} \right. \\
&\quad \left. \times \left( p_i \times \frac{e^{-\theta_i\tau_j}(\theta_i\tau_j)^{y_{ij}}}{y_{ij}!} \right)^{z_{ij}I(y_{ij}>0)} \right] \\
&\quad \times f(p_i|a_1, b_1) \times f(\theta_i|\alpha_i, \beta_i) \times f(\alpha_i|a_2, b_2) \times f(\beta_i|a_3, b_3) \\
&= \left[ \prod_{j=1}^{n} \left( [(1-p_i)\delta(0)]^{1-z_{ij}} \right)^{I(y_{ij}=0)} \times p_i^{z_{ij}} \left[ \frac{e^{-\theta_i\tau_j}(\theta_i\tau_j)^{y_{ij}}}{y_{ij}!} \right]^{z_{ij}} \right. \\
&\quad \times f(p_i|a_1, b_1) \times f(\theta_i|\alpha_i, \beta_i)] \times f(\alpha_i|a_2, b_2) \times f(\beta_i|a_3, b_3) \\
&= \left[ \prod_{j=1}^{n} \left( [(1-p_i)\delta(0)]^{1-z_{ij}} \right)^{I(y_{ij}=0)} \times p_i^{z_{ij}} \left[ \frac{e^{-\theta_i\tau_j}(\theta_i\tau_j)^{y_{ij}}}{y_{ij}!} \right]^{z_{ij}} \right. \\
&\quad \left. \times \frac{p_i^{a_1-1} \times (1-p_i)^{b_1-1}}{B(a_1, b_1)} \times \frac{\beta_i^{\alpha_i}}{\Gamma(\alpha_i)} \theta_i^{\alpha_i-1} e^{-\beta_i\theta_i} \right] \\
&\quad \times \frac{b_2^{a_2}}{\Gamma(a_2)} \alpha_i^{a_2-1} e^{-b_2\alpha_i} \times \frac{b_3^{a_3}}{\Gamma(a_3)} \beta_i^{a_3-1} e^{-b_3\beta_i}
\end{aligned}
$$

Posterior distributions are derived as below:

$$
\begin{aligned}
f(\theta_i|p_i, \alpha_i, \beta_i, \boldsymbol{y}_i, \boldsymbol{z}_i, \tau_j) &\propto \theta_i^{\alpha_i + \sum_{j=1}^{n} y_{ij}z_{ij} - 1} e^{-(\beta_i + \sum_{j=1}^{n} \tau_j z_{ij})\theta_i} \\
&\propto \text{Gamma}(\alpha_i + \sum_{j=1}^{n} y_{ij}z_{ij}, \beta_i + \sum_{j=1}^{n} \tau_j z_{ij})
\end{aligned}
$$

$$
\begin{aligned}
f(p_i|\theta_i, \alpha_i, \beta_i, \boldsymbol{y}_i, \boldsymbol{z}_i, \tau_j) &\propto \prod_{j=1}^{n} p_i^{z_{ij}} (1-p_i)^{(1-z_{ij})I(y_{ij}=0)} \times p_i^{a_1-1} \times (1-p_i)^{b_1-1} \\
&\propto \text{Beta}(a_1 + \sum_{j=1}^{n} z_{ij}, b_1 + \sum_{j=1}^{n} (1-z_{ij})I(y_{ij}=0))
\end{aligned}
$$

$$
f(\alpha_i|\theta_i, p_i, \beta_i, \boldsymbol{y}_i, \boldsymbol{z}_i, \tau_j) \propto \frac{\alpha_i^{a_2-1}}{\Gamma(\alpha_i)} \times e^{\alpha_i(-b_2 + \cdot log(\beta_i) + log(\theta_i))}
$$

We follow the random walk proposal from Jia et al. (2017) to simulate the conditional distribution $f(\alpha_i|\cdot)$.

$$f(\beta_i|\theta_i, p_i, \alpha_i, \boldsymbol{y}_i, \boldsymbol{z}_i, \tau_j) \propto \beta_i^{a_3+\alpha_i-1} \times e^{-\beta_i(b_3+\theta_i)}$$
$$\propto \text{Gamma}(a_3+\alpha_i, b_3+\theta_i)$$

The details of the derivation of the posterior distribution for $z_{ij}$ is as follows.

$$P(z_{ij}=1|y_{ij},\theta_i,p_i,\alpha_i,\beta_i,\tau_j) = \frac{P(z_{ij}=1,y_{ij},\theta_i,p_i,\alpha_i,\beta_i,\tau_j)}{P(z_{ij}=1,y_{ij},\theta_i,p_i,\alpha_i,\beta_i,\tau_j)+P(z_{ij}=0,y_{ij},\theta_i,p_i,\alpha_i,\beta_i,\tau_j)}$$
$$= \frac{p_i e^{-\theta_i\tau_j}(\theta_i\tau_j)^{y_{ij}}/y_{ij}!}{p_i e^{-\theta_i\tau_j}(\theta_i\tau_j)^{y_{ij}}/y_{ij}! + [(1-p_i)\delta(0)]^{I(y_{ij}=0)}}$$

The posterior distribution for $z_{ij}$ is:

$$f(z_{ij}|\theta_i,p_i,\alpha_i,\beta_i,\boldsymbol{y}_i,\tau_j) \propto \text{Bernoulli}\left(\frac{p_i e^{-\theta_i\tau_j}(\theta_i\tau_j)^{y_{ij}}/y_{ij}!}{p_i e^{-\theta_i\tau_j}(\theta_i\tau_j)^{y_{ij}}/y_{ij}! + [(1-p_i)\delta(0)]^{I(y_{ij}=0)}}\right)$$
$$\propto \text{Bernoulli}\left(\frac{p_i e^{-\theta_i\tau_j}}{p_i e^{-\theta_i\tau_j} + (1-p_i)I(y_{ij}=0)}\right)$$

Following the Lemma 1 and Lemma 2 from Jia et al. (2017), we set $a_2, a_3$ to small positive values, set $b_2, b_3$ to large numbers, and increase the values of $b_2, b_3$ in iteration $t$ by:

$$b_2^{(t)} = b_2^{(t-1)} + \frac{1}{t}, \quad b_3^{(t)} = b_3^{(t-1)} + \frac{1}{t}.$$

Thus, it can be shown that the posterior mean $E(\alpha_i|\theta_i,p_i,\beta_i,\boldsymbol{y}_i,\boldsymbol{z}_i,\tau_j) \to 0$ if $b_2 \to \infty$; $E(\beta_i|\theta_i,p_i,\alpha_i,\boldsymbol{y}_i,\boldsymbol{z}_i,\tau_j) \to 0$ as $b_3 \to \infty$.

For gene $i$, let $n_{0i} = \sum_{j=1}^n (1-z_{ij})$ and $n_{1i} = \sum_{j=1}^n z_{ij}$, $n = n_{1i} + n_{0i}$. If $\frac{a_1}{a_1+b_1} \approx \frac{n_{1i}}{n}$, then we have $E(p_i|\theta_i,\alpha_i,\beta_i,\boldsymbol{y}_i,\boldsymbol{z}_i,\tau_j) = \frac{a_1+\sum_{j=1}^n z_{ij}}{a_1+\sum_{j=1}^n z_{ij}+b_1+\sum_{j=1}^n (1-z_{ij})I(y_{ij}=0)} = \frac{a_1+n_{1i}}{a_1+b_1+n_i} \approx \frac{n_{1i}}{n_i}$. Therefore, the values $a_1$ and $b_1$ should be chosen based on an estimate of the dropout rate of the dataset. Ideally, $\frac{a_1}{b_1} = \frac{n_{1i}}{n_{0i}}$. For simplicity, $\frac{a_1}{b_1} = median(\frac{n_{1i}}{n_{0i}})$.

$E(\theta_i|p_i,\alpha_i,\beta_i,\boldsymbol{y}_i,\boldsymbol{z}_i,\tau_j) = \frac{\alpha_i+\sum_{j=1}^n y_{ij}z_{ij}}{\beta_i+\sum_{j=1}^n \tau_j z_{ij}} \to \frac{\sum_{j=1}^n y_{ij}}{\sum_{j=1}^n \tau_j z_{ij}}$.

$E(z_{ij}|\theta_i,p_i,\alpha_i,\beta_i,\boldsymbol{y}_i,\tau_j) = \frac{p_i e^{-\theta_i\tau_j}}{p_i e^{-\theta_i\tau_j}+(1-p_i)I(y_{ij}=0)}$.

#### 2. Generation of the covariance and precision matrices with known structure

This section largely follows Lyu et al. (2018).

(i) Split genes into $L$ blocks. Genes within each block are more likely to be correlated.

(ii) For each block $l$, simulate a square matrix whose elements are generated as $A^{(l)}(i,i')$.

$$A^{(l)}(i,i') = \begin{cases} 1 & \text{if } i = i',\ i = 1, ..., G \\ 0 & \text{if } i \neq i', \text{ gene } i \text{ and gene } i' \text{ are not conditionally correlated} \\ \sim U(D) & \text{gene } i \text{ and gene } i' \text{ are conditionally correlated} \end{cases} \quad (1)$$

where $U(D)$ is a uniform distribution with $D = [-1, -0.6] \cup [0.6, 1]$. A diagonal matrix is then added to $A^{(l)}$ to ensure that the derived matrix is positive definite: $B^{(l)} = A^{(l)} + \delta I$. $\delta$ is a selected constant.

(iii) Calculate the matrix $\Sigma^{(l)}(i,i') = [B^{(l)}]^{-1}(i,i')/\sqrt{[B^{(l)}]^{-1}(i,i)[B^{(l)}]^{-1}(i',i')}$ for block $l,\ l = 1, 2, ..., L$.

(iv) Obtain the overall covariance matrix $\Sigma = \begin{bmatrix} \Sigma^{(1)} & & & \\ & \Sigma^{(2)} & & \\ & & \cdots & \\ & & & \Sigma^{(L)} \end{bmatrix}$.

(v) The matrix $\Sigma$ will serve as the covariance matrix of the multivariate Gaussian distribution in step (a) in the simulation session. The inverse matrix of $\Sigma$ is the desired precision matrix $\Omega = \Sigma^{-1}$.

#### 3. Adapted StARS method in JFGL

Following Liu et al. (2010), we adapt the StARS method to the joint network scenario as follows:

(1) Fix $\lambda_1$ value, choose a set of candidate values $\{\lambda_2^{(1)}, ..., \lambda_2^{(L)}\}$ for $\lambda_2$. $L$ is the total

number of candidate values. The initial value of $\lambda_1$ could be a small constant such as 0.05.

(2) Calculate subsampling ratio based on the sample size (Zhao et al., 2012) under each condition $k$: the ratio is 0.8 if $n_k \leqslant 144$; the ratio is $10 \times \sqrt{n_k}/n_k$ if $n_k > 144$

(3) Take subsamples from the original data according to the subsampling ratio calculated in step (2). Fit JGL model using the fixed $\lambda_1$ value and each of the candidate $\lambda_2$ values separately. For each fitted model, we can derive the estimated precision matrices $\{\hat{\Omega}^{(1)}_{\lambda_1,\lambda_2^{(l)}}, ..., \hat{\Omega}^{(K)}_{\lambda_1,\lambda_2^{(l)}}\}$, $l = 1, 2, ..., L$. Dichotomize the estimated precision matrices by forcing the non-zero values to ones, and thus we derive the adjacency matrices $\{\hat{Adj}^{(1)}_{\lambda_1,\lambda_2^{(l)}}, ..., \hat{Adj}^{(K)}_{\lambda_1,\lambda_2^{(l)}}\}$, $l = 1, 2, ..., L$.

(4) Repeat step (3) for $n_{rep}$ times. For each $\lambda_2^{(l)}$, calculate the empirical edge selection probabilities under each condition (denote the matrix by $P^{(k)}_{\lambda_1,\lambda_2^{(l)}}$) by taking the average of the sum of the $n_{rep}$ adjacency matrices.

(5) For each $\lambda_2^{(l)}$, calculate the sum of edge variabilities for the joint model by taking the sum of elements from the matrix: $\sum_{k=1}^{K} 4 \times \left[P^{(k)}_{\lambda_1,\lambda_2^{(l)}} \circ (1 - P^{(k)}_{\lambda_1,\lambda_2^{(l)}})\right] / (G \times (G-1))$, where $\circ$ denotes the element-wise multiplication between matrices.

(6) Select the optimal $\lambda_2$ value by taking the supremum edge variability value among the edge variabilities that are less than the supplied threshold $t_s$. The standard $t_s$ value is 0.05 or 0.1 (Liu et al., 2010; Kurtz et al., 2015; Yoon et al., 2019)

(7) To derive the optimal $\lambda_1$ value, fix the optimal $\lambda_2$ value and repeat similar steps as the above (1) to (4) using a set of candidate values $\{\lambda_1^{(1)}, ..., \lambda_1^{(L)}\}$ for $\lambda_1$.

#### 4. Real Data Processing Details

The R package Seurat (Butler et al., 2018) is used for single cell clustering for both Medulloblastoma (Hovestadt et al., 2019) and Glioblastoma (Neftel et al., 2019) samples. To create Seurat objects, TPM count matrices are used as input. Genes expressed in at least 3 cells are kept; cells expressed at least 200 genes are kept. "LogNormalize" with scale factor 10000 is used to normalize the filtered data. The top 2000 variable genes are selected and used for the dimension reduction steps.

For the purpose of joint network construction, we take the gene subset where at least 20 cells have the gene expression $> 0$ in each group. For the JGNsc MCMC continuization step, the burn-in number is set to 500 and the iteration number is set to 500. AIC is used to select the tuning parameter in the JGL model.

#### 5. Supplemental Figures

[Figure 1 about here.]

[Figure 2 about here.]

[Figure 3 about here.]

[Figure 4 about here.]

[Figure 5 about here.]

[Figure 6 about here.]

[Figure 7 about here.]

[Figure 8 about here.]

[Figure 9 about here.]

[Figure 10 about here.]

[Figure 11 about here.]

[Figure 12 about here.]

[Figure 13 about here.]

[Figure 14 about here.]

[Figure 15 about here.]

#### References

Butler, A., Hoffman, P., Smibert, P., Papalexi, E., and Satija, R. (2018). Integrating single-cell transcriptomic data across different conditions, technologies, and species. *Nature biotechnology* **36,** 411–420.

Hovestadt, V., Smith, K. S., Bihannic, L., Filbin, M. G., Shaw, M. L., Baumgartner, A., DeWitt, J. C., Groves, A., Mayr, L., Weisman, H. R., et al. (2019). Resolving medulloblastoma cellular architecture by single-cell genomics. *Nature* **572,** 74–79.

Jia, B., Xu, S., Xiao, G., Lamba, V., and Liang, F. (2017). Learning gene regulatory networks from next generation sequencing data. *Biometrics* **73,** 1221–1230.

Kurtz, Z. D., Müller, C. L., Miraldi, E. R., Littman, D. R., Blaser, M. J., and Bonneau, R. A. (2015). Sparse and compositionally robust inference of microbial ecological networks. *PLoS Comput Biol* **11,** e1004226.

Liu, H., Roeder, K., and Wasserman, L. (2010). Stability approach to regularization selection (stars) for high dimensional graphical models. In *Advances in neural information processing systems*, pages 1432–1440.

Lyu, Y., Xue, L., Zhang, F., Koch, H., Saba, L., Kechris, K., and Li, Q. (2018). Condition-adaptive fused graphical lasso (cfgl): An adaptive procedure for inferring condition-specific gene co-expression network. *PLoS computational biology* **14,** e1006436.

Neftel, C., Laffy, J., Filbin, M. G., Hara, T., Shore, M. E., Rahme, G. J., Richman, A. R., Silverbush, D., Shaw, M. L., Hebert, C. M., et al. (2019). An integrative model of cellular states, plasticity, and genetics for glioblastoma. *Cell* **178,** 835–849.

Yoon, G., Gaynanova, I., and Müller, C. L. (2019). Microbial networks in spring-semi-parametric rank-based correlation and partial correlation estimation for quantitative microbiome data. *Frontiers in genetics* **10,** 516.

Zhao, T., Liu, H., Roeder, K., Lafferty, J., and Wasserman, L. (2012). The huge package for high-dimensional undirected graph estimation in r. *Journal of Machine Learning Research* **13,** 1059–1062.

*Received xxx* 202x. *Revised xxx* 2021. *Accepted xxx* 2021.

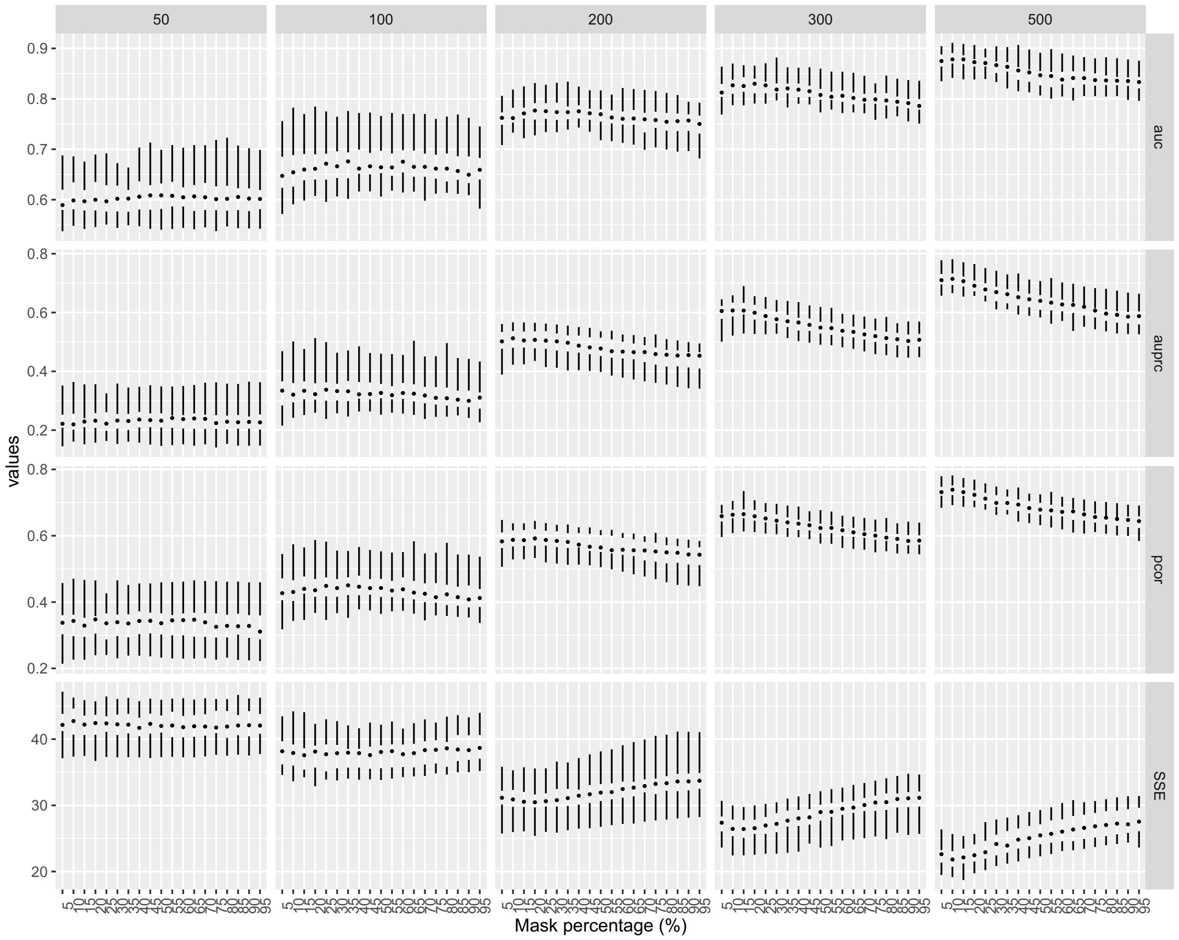


**Figure 1.** JGNsc_Hybrid mask rate benchmark. The mask rate varies from 5% to 95%. From the simulation, 10% $\sim$ 20% of mask rate achieves better performance. 15% is used as the mask rate in the methods benchmark sections.

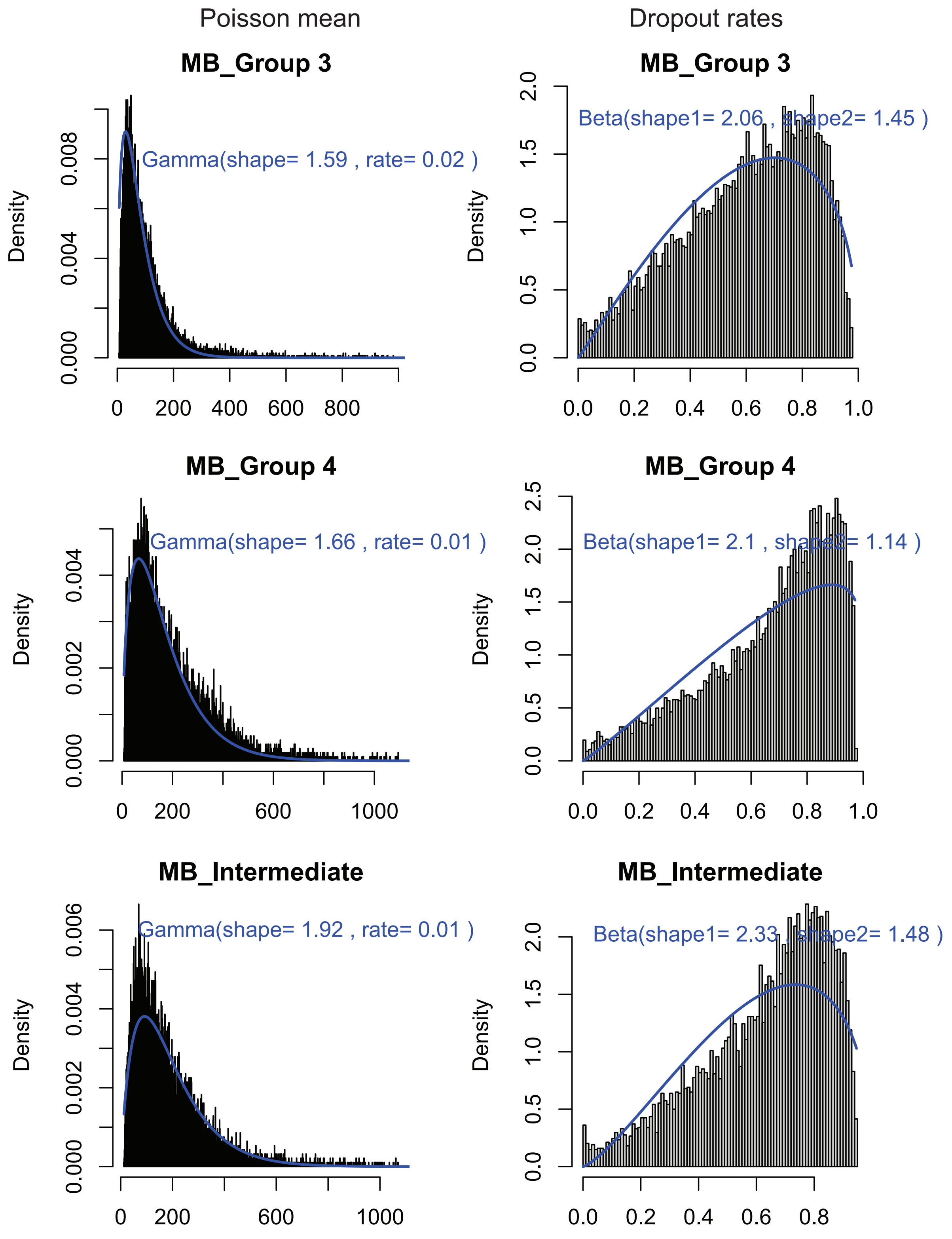


**Figure 2.** Estimated distribution of hyperparameters using the MB data. R package GAMLSS is applied.

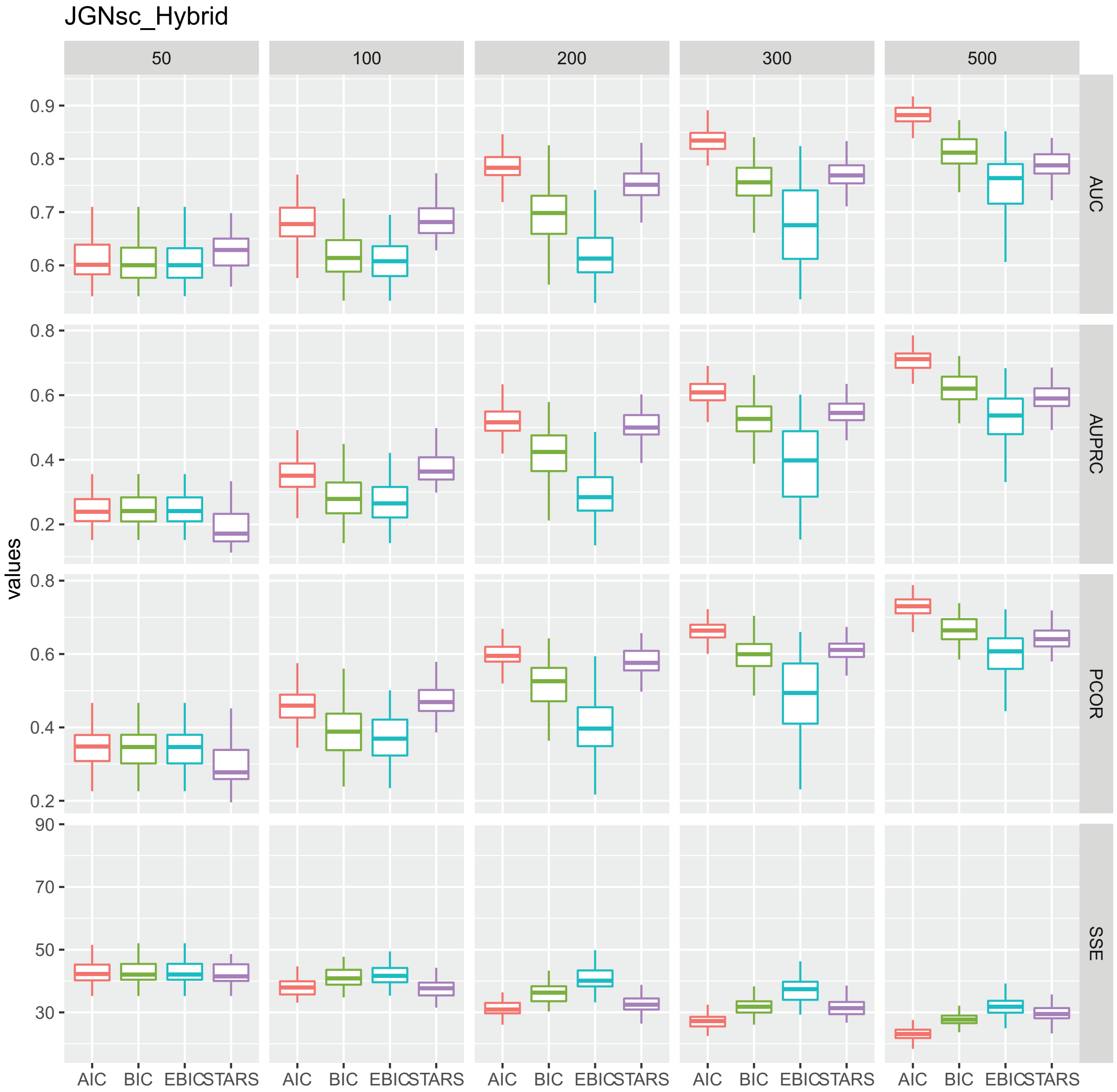


**Figure 3.** Tuning parameter benchmark for JGNsc_Hybrid by Joint Graphical Lasso. Scenario: partially identical precision matrix structures – the first 20 genes having different precision structures and weights. AIC outperforms the other three criteria in all four metrics when sample size is greater than the number of genes (100). AIC and StARS perform similar when the sample size is less than or equal to the number of genes. AUC: Area under ROC curve; AUPRC: Area under precision recall curve; PCOR: pearson correlation; SSE: sum of squared errors.

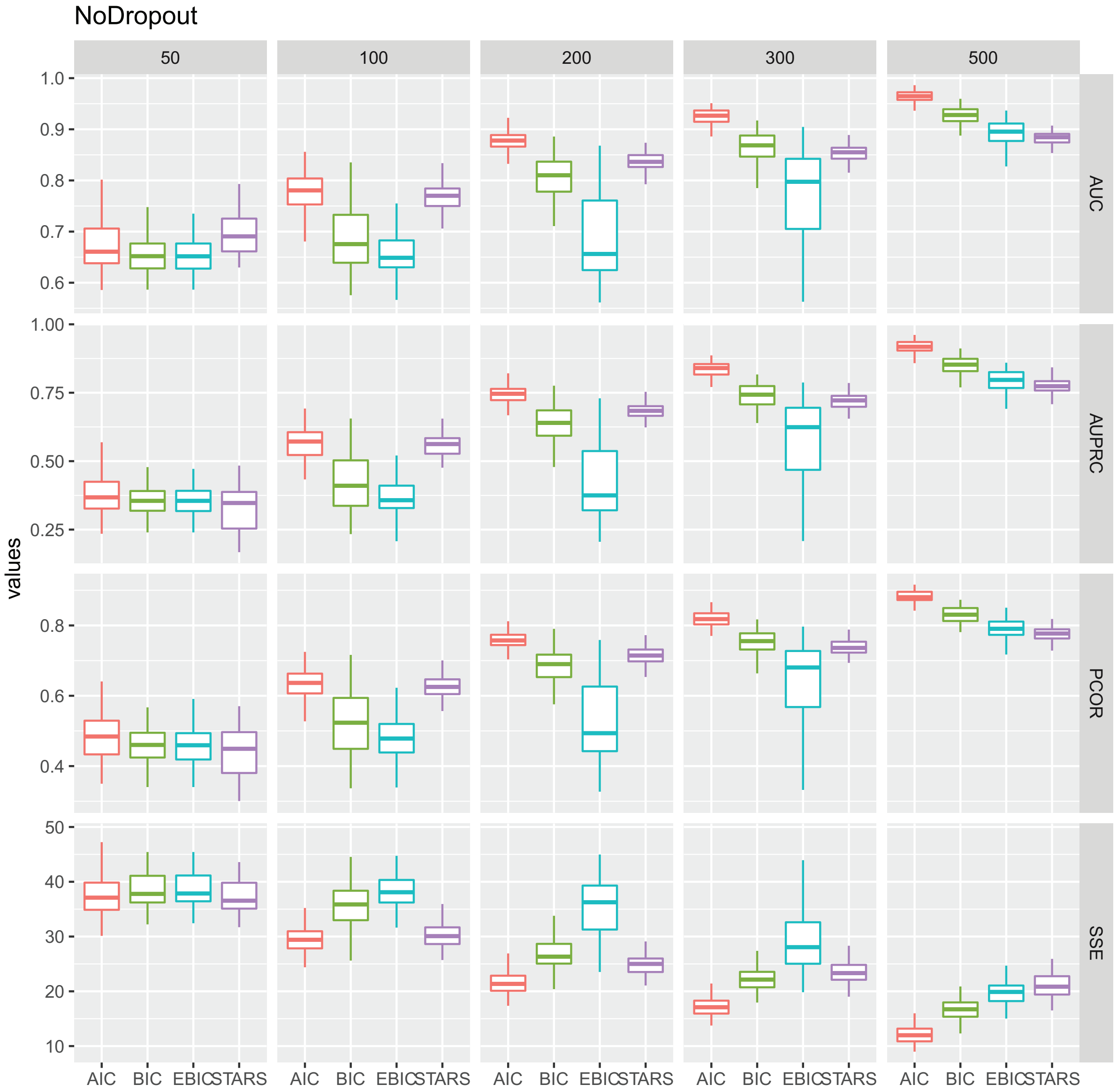


**Figure 4.** Tuning parameter benchmark for No Dropout scenario by Joint Graphical Lasso. Scenario: partially identical precision matrix structures – the first 20 genes having different precision structures and weights. AIC outperforms the other three criteria in all four metrics. AUC: Area under ROC curve; AUPRC: Area under precision recall curve; PCOR: pearson correlation; SSE: sum of squared errors.

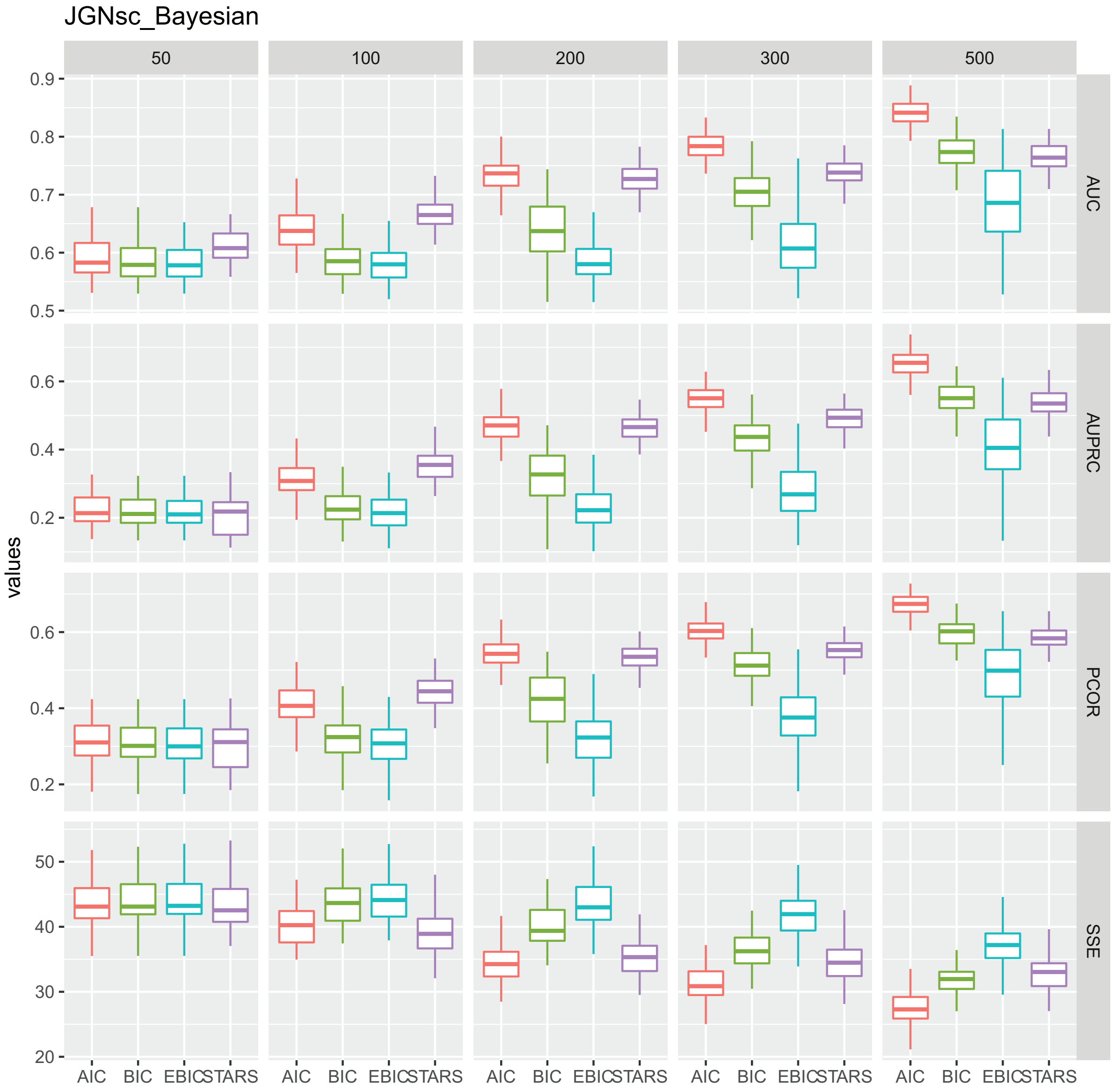


**Figure 5.** Tuning parameter benchmark for JGNsc by Joint Graphical Lasso. Scenario: partially identical precision matrix structures – the first 20 genes having different precision structures and weights. AIC outperforms the other three criteria in all four metrics when sample size is greater than the number of genes (100). StARS performs slightly better than AIC when the sample size is less than or equal to the number of genes. AUC: Area under ROC curve; AUPRC: Area under precision recall curve; PCOR: pearson correlation; SSE: sum of squared errors.

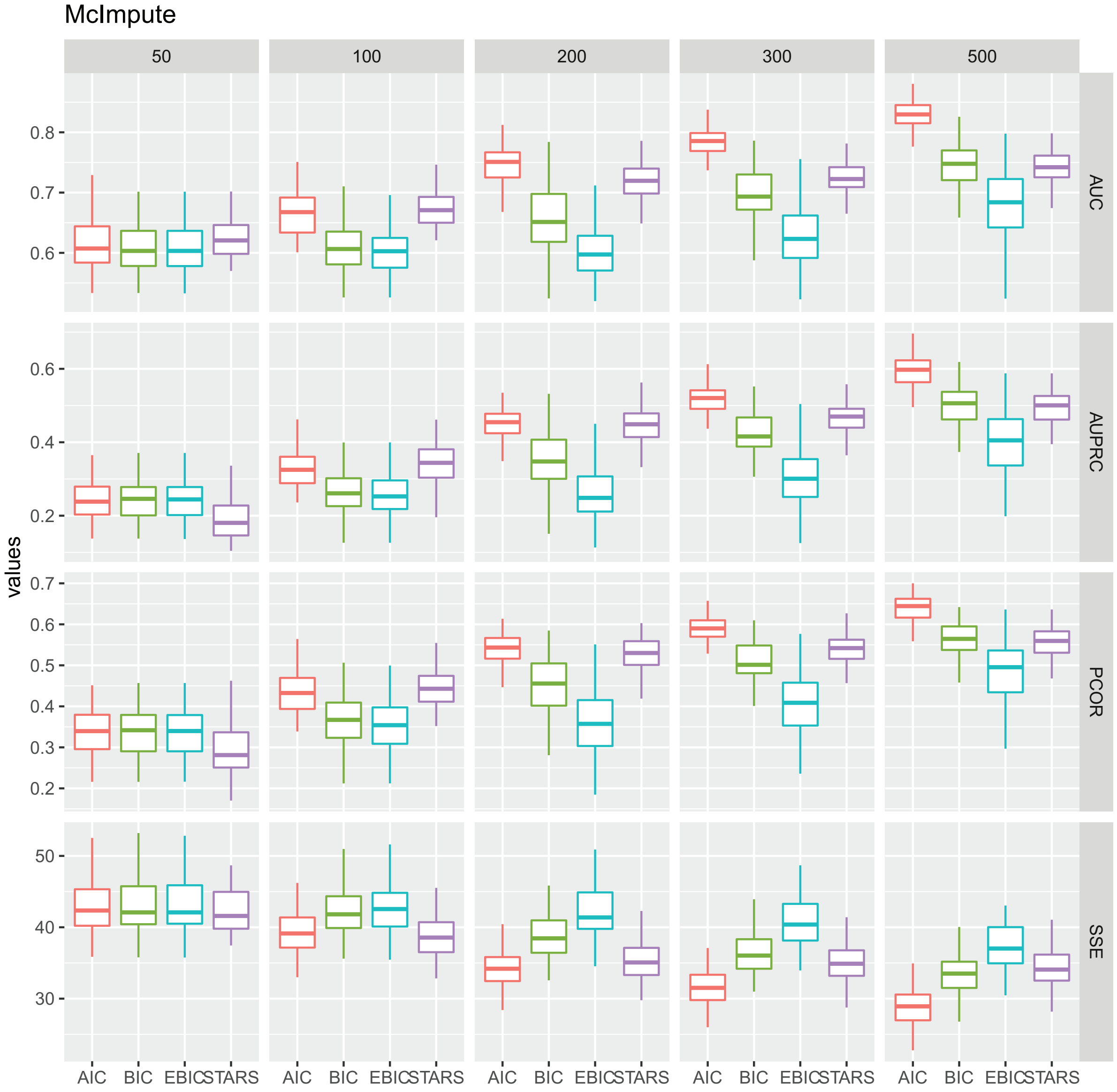


**Figure 6.** Tuning parameter benchmark for McImpute by Joint Graphical Lasso. Scenario: partially identical precision matrix structures – the first 20 genes having different precision structures and weights. AIC outperforms the other three criteria in all four metrics when sample size is greater than the number of genes (100). AIC and StARS perform similar when the sample size is less than or equal to the number of genes. AUC: Area under ROC curve; AUPRC: Area under precision recall curve; PCOR: pearson correlation; SSE: sum of squared errors.

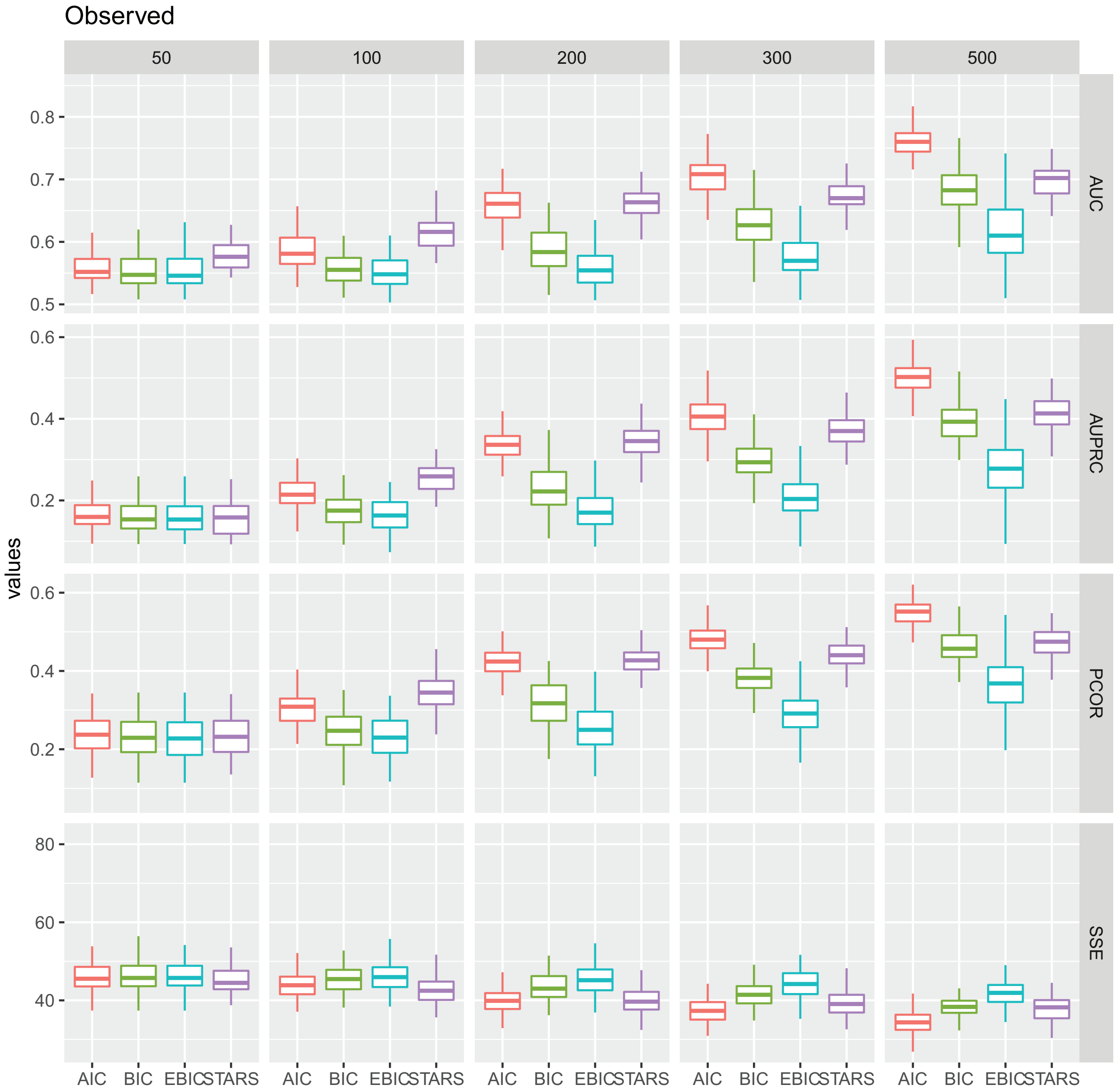


**Figure 7.** Tuning parameter benchmark for Observed counts by Joint Graphical Lasso. AIC outperforms the other three criteria in all four metrics when sample size is greater than 200. StARS performs slightly better than AIC when the sample size is less than or equal to 200. AUC: Area under ROC curve; AUPRC: Area under precision recall curve; PCOR: pearson correlation; SSE: sum of squared errors.

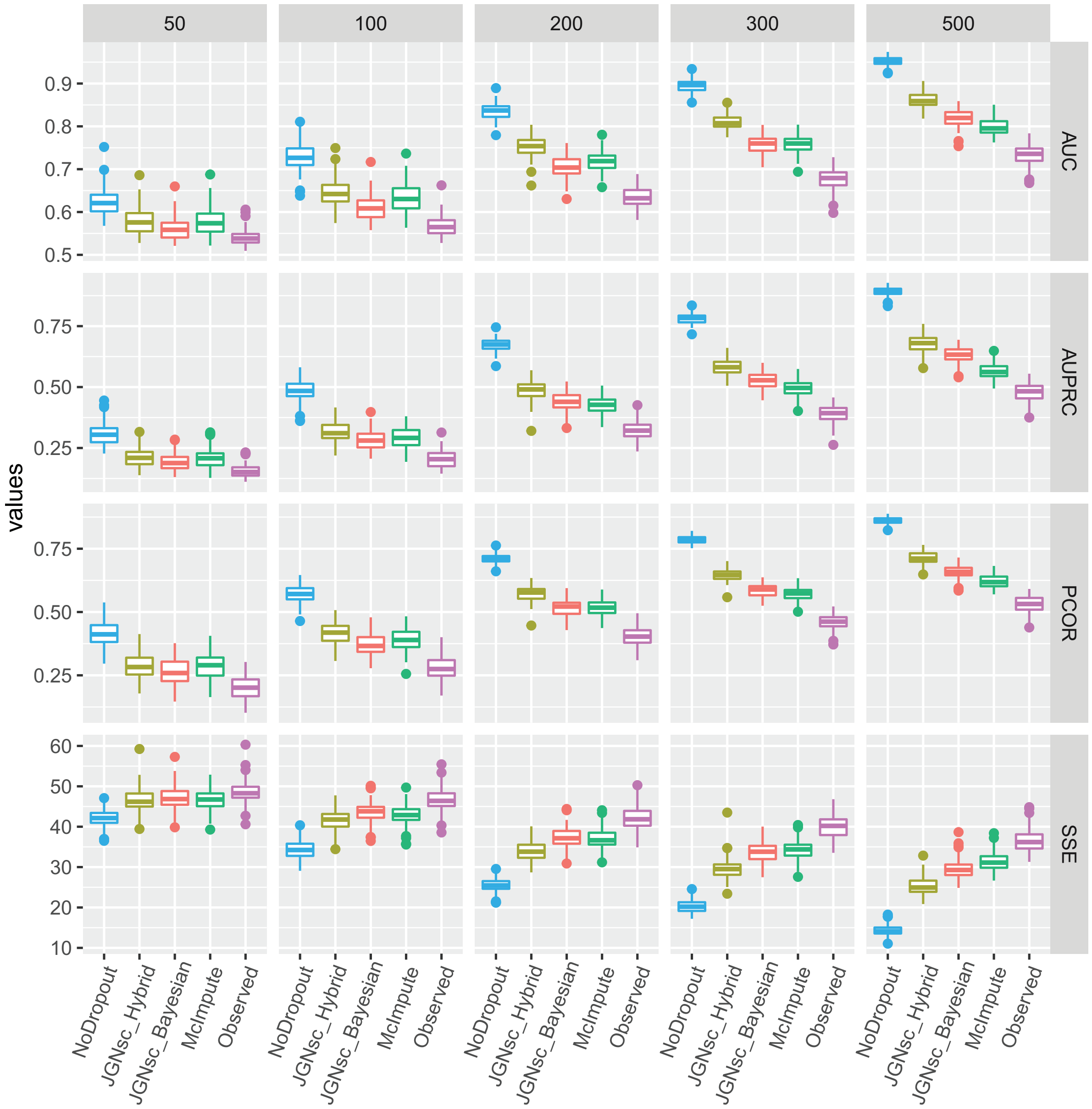


**Figure 8.** Benchmark data processing methods, scenario: partially identical precision matrix structures – the first 50 genes having different precision structures and weights. JGNsc_Hybrid outperforms JGNsc, McImpute and Observed scenario as sample size varies. The performance of JGNsc without iteration becomes better and excel McImpute when the sample size is large (500).

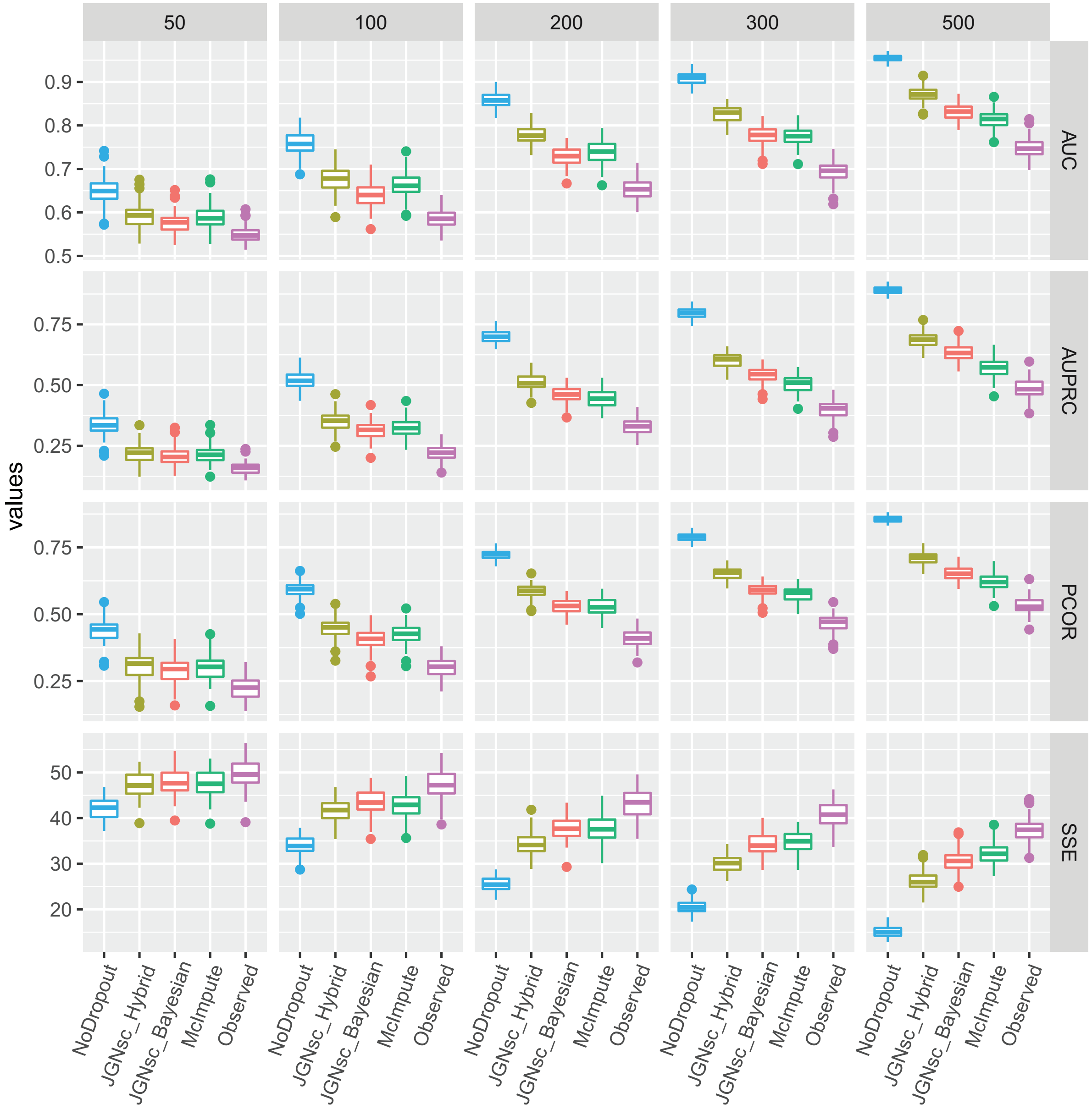


**Figure 9.** Benchmark data processing methods, scenario: identical precision matrix structures with different weights. JGNsc_Hybrid outperforms JGNsc, McImpute and Observed scenario as sample size varies. The performance of JGNsc without iteration becomes better and excel McImpute when the sample size is large (500).

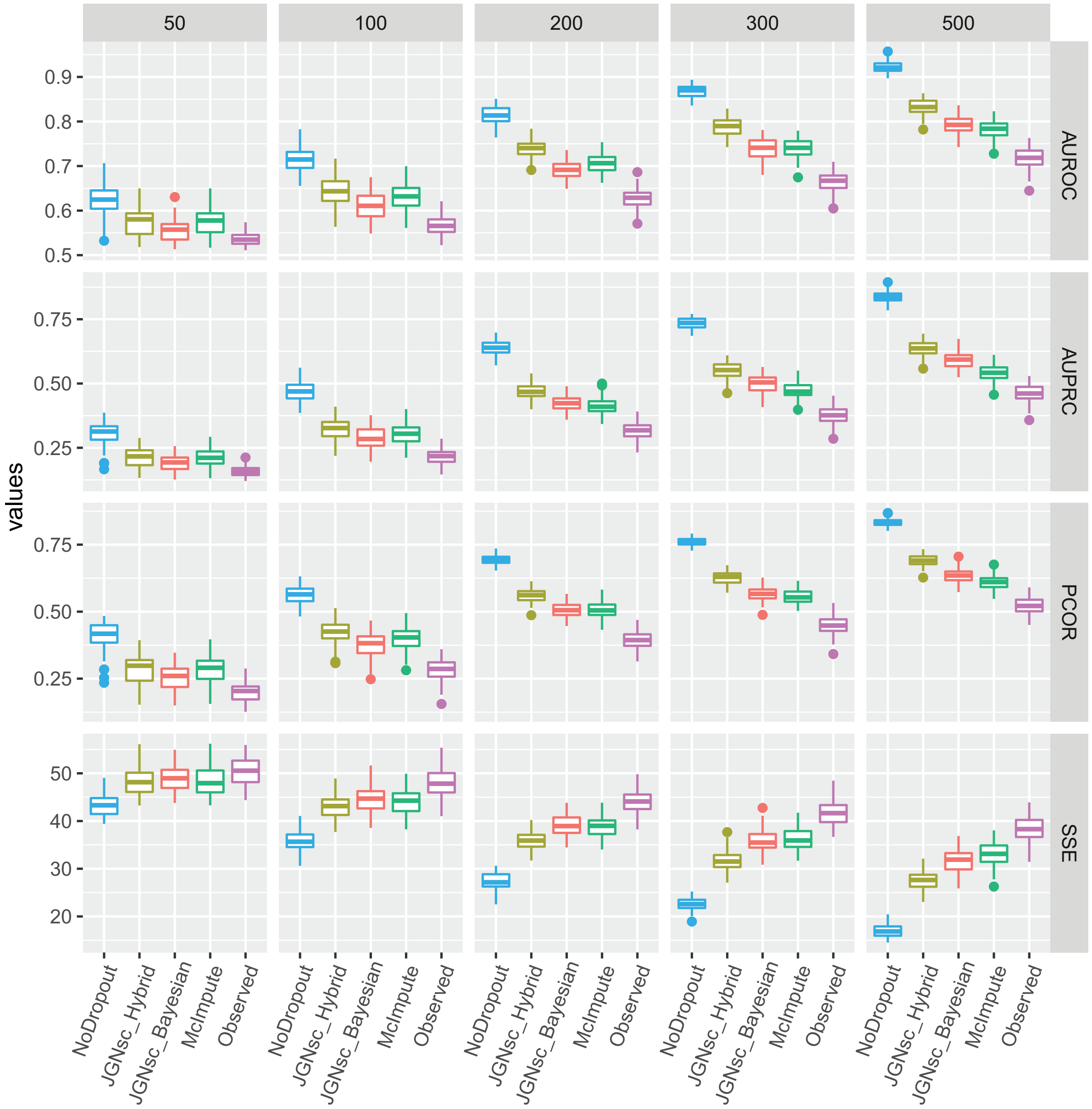


**Figure 10.** Benchmark data processing methods, scenario: different precision matrix structures and weights. JGNsc_Hybrid outperforms JGNsc, McImpute and Observed scenario as sample size varies. The performance of JGNsc without iteration becomes better and excel McImpute when the sample size is large (500).

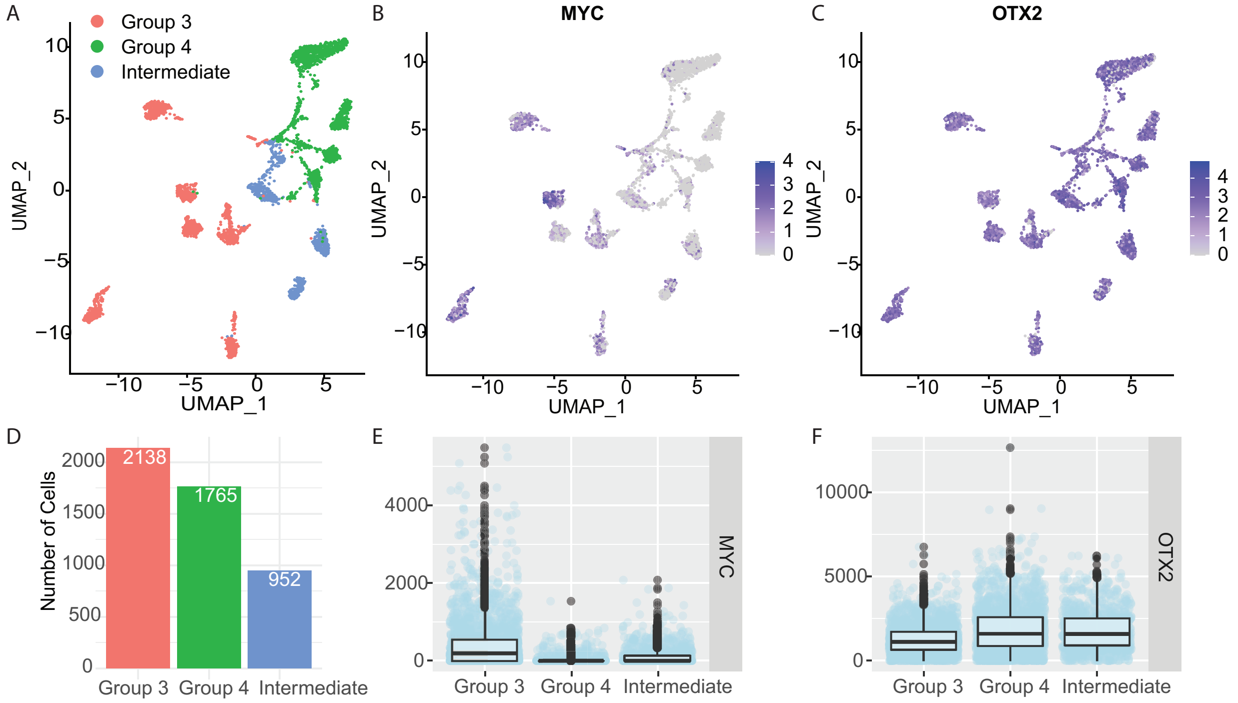


**Figure 11.** Demographic information of MB data. (A) UMAP plot of scRNA-seq data from 17 selected patients from Hovestadt et al. (2019). The samples were classified into Group 3, Group 4 and the intermediate group by DNA mythelation. As pointed out in the original paper, MYC is an important biomarker for Medulloblastoma, and is associated with unfavorable results. OTX2 is a protein coding gene that is closely related to brain development. Therefore (B) and (C) shows their expression levels in a UMAP plot. The color shows the normalized counts from Seurat. The feature expression measurements for each cell are divided by the total expression, then multiplied by a scale factor 10,000, and log-transformed. (D) The number of single cells in each group. (E) and (F) are the boxplot that summarize the expression levels of MYC and OTX2. For MYC, we observe the non-zero expression rates are $74\%, 9\%, 38\%$ for Group 3, Group 4 and Intermediate group. For OTX2, we observe the non-zero expression rates are $98\%, 97\%, 98\%$ for Group 3, Group 4 and Intermediate group.

A

| pathways | *Metabolism enzymes gene set* | Composition of genes connected to **MYC** | | Fisher's exact test p-value | |
|---|---|---|---|---|---|
| | | Group 3 | Intermediate group | Group 3 | Intermediate group |
| Anaplerotic Reactions of the TCA Cycle | 0.02 | 0.07 | 0.00 | 0.01 | 1.00 |
| Deamination of Amino Acids: The Urea Cycle | 0.03 | 0.03 | 0.00 | 0.69 | 1.00 |
| Folic Acid Metabolism | 0.02 | 0.05 | 0.00 | 0.17 | 1.00 |
| Folic Acid Reactions | 0.01 | 0.03 | 0.00 | 0.16 | 1.00 |
| Gluconeogenesis and Glycolysis | 0.05 | 0.12 | 0.17 | 0.03 | 0.28 |
| Gluconeogenesis and Glycolysis: Regulation | 0.02 | 0.03 | 0.00 | 0.26 | 1.00 |
| Glycolysis and TCA Cycle | 0.06 | 0.15 | 0.17 | <0.01 | 0.30 |
| Heme Metabolism | 0.02 | 0.00 | 0.00 | 1.00 | 1.00 |
| Isoprene Metabolism | 0.07 | 0.05 | 0.00 | 0.79 | 1.00 |
| Lipid Metabolism | 0.15 | 0.17 | 0.33 | 0.70 | 0.22 |
| Pentose Phosphate Pathways | 0.02 | 0.02 | 0.00 | 1.00 | 1.00 |
| Products of Isoprene Metabolism | 0.09 | 0.08 | 0.17 | 1.00 | 0.42 |
| Prostaglandins, Thromboxanes, and Leucotrienes | 0.02 | 0.02 | 0.00 | 1.00 | 1.00 |
| Purine Metabolism | 0.17 | 0.22 | 0.17 | 0.28 | 1.00 |
| Pyrimidine Metabolism | 0.09 | 0.08 | 0.00 | 1.00 | 1.00 |
| Undefined | 0.42 | 0.41 | 0.33 | 0.89 | 1.00 |
| Total number of genes connected to MYC | (Total: 651) | 59 | 6 | | |

B

| pathways | *Metabolism enzymes gene set* | Composition of genes connected to **OTX2** | | | Fisher's exact test p-value | | |
|---|---|---|---|---|---|---|---|
| | | Group 3 | Intermediate group | Group 4 | Group 3 | Intermediate group | Group 4 |
| Anaplerotic Reactions of the TCA Cycle | 0.02 | 0.04 | 0.03 | 0.14 | 0.32 | 0.38 | 0.10 |
| Deamination of Amino Acids: The Urea Cycle | 0.03 | 0.04 | 0.00 | 0.14 | 0.52 | 1.00 | 0.19 |
| Folic Acid Metabolism | 0.02 | 0.00 | 0.00 | 0.00 | 1.00 | 1.00 | 1.00 |
| Folic Acid Reactions | 0.01 | 0.00 | 0.00 | 0.00 | 1.00 | 1.00 | 1.00 |
| Gluconeogenesis and Glycolysis | 0.05 | 0.08 | 0.17 | 0.00 | 0.36 | 0.02 | 0.39 |
| Gluconeogenesis and Glycolysis: Regulation | 0.02 | 0.04 | 0.07 | 0.00 | 0.34 | 0.09 | 1.00 |
| Glycolysis and TCA Cycle | 0.06 | 0.04 | 0.20 | 0.00 | 1.00 | 0.00 | 0.41 |
| Heme Metabolism | 0.02 | 0.00 | 0.03 | 0.00 | 1.00 | 0.49 | 1.00 |
| Isoprene Metabolism | 0.07 | 0.04 | 0.13 | 0.00 | 1.00 | 0.14 | 0.25 |
| Lipid Metabolism | 0.15 | 0.29 | 0.10 | 0.29 | 0.07 | 0.60 | 0.28 |
| Pentose Phosphate Pathways | 0.02 | 0.00 | 0.10 | 0.00 | 1.00 | 0.02 | 1.00 |
| Products of Isoprene Metabolism | 0.09 | 0.08 | 0.20 | 0.00 | 1.00 | 0.04 | 0.10 |
| Prostaglandins, Thromboxanes, and Leucotrienes | 0.02 | 0.00 | 0.00 | 0.00 | 1.00 | 1.00 | 1.00 |
| Purine Metabolism | 0.17 | 0.17 | 0.07 | 0.14 | 1.00 | 0.21 | 1.00 |
| Pyrimidine Metabolism | 0.09 | 0.08 | 0.07 | 0.14 | 1.00 | 1.00 | 0.48 |
| Undefined | 0.42 | 0.38 | 0.47 | 0.43 | 0.83 | 0.71 | 1.00 |
| Total number of genes connected to MYC | (Total: 651) | 24 | 30 | 7 | | | |

**Figure 12.** Gene set enrichment analysis result for MB data. To learn functional categories of the genes connected with (A) MYC (B) OTX2, these genes are mapped to the NCBI BioSystems Database pathways.

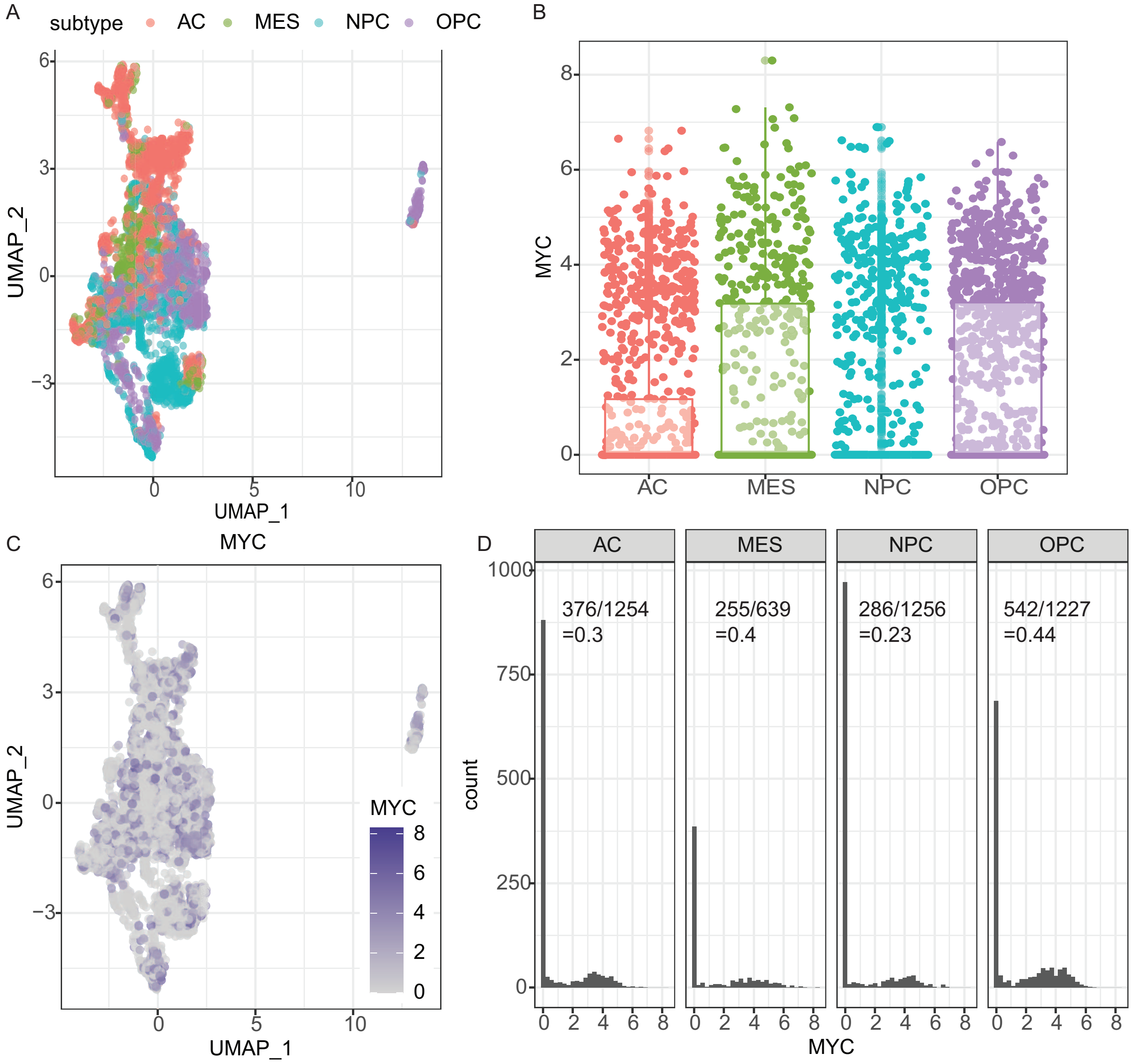


**Figure 13.** GBM scRNA-seq data characteristics from Neftel et al. (2019). (A) UMAP plot of the clustering result from Seurat (Butler et al., 2018) of the selected subset of single cell samples. Samples are classified into four groups: AC-like, MES-like, NPC-like and OPC-like cells. (B) Boxplot overlaid with jitter plot of the expression level of MYC in each sub-group. Y-axis is the normalized counts from Seurat. The feature expression measurements for each cell are divided by the total expression, then multiplied by a scale factor 10,000, and log-transformed. Along with figure (D) below, we see the median of MYC expression level in all four groups are at zero. The upper edge of the "boxplot" shows the 75% quantile of the data. For example, we observe that the majority of NPC-like cells have zero MYC expression. Hence, with only 23% of non-zero counts, the boxplot of NPC-like cells overlapped with the zero line. (C) Feature plot of the gene expression of MYC. (D) Histogram of the distribution of MYC expression levels. The number in each panel shows the fraction of cells that have MYC expressed ($count > 0$) and captured.

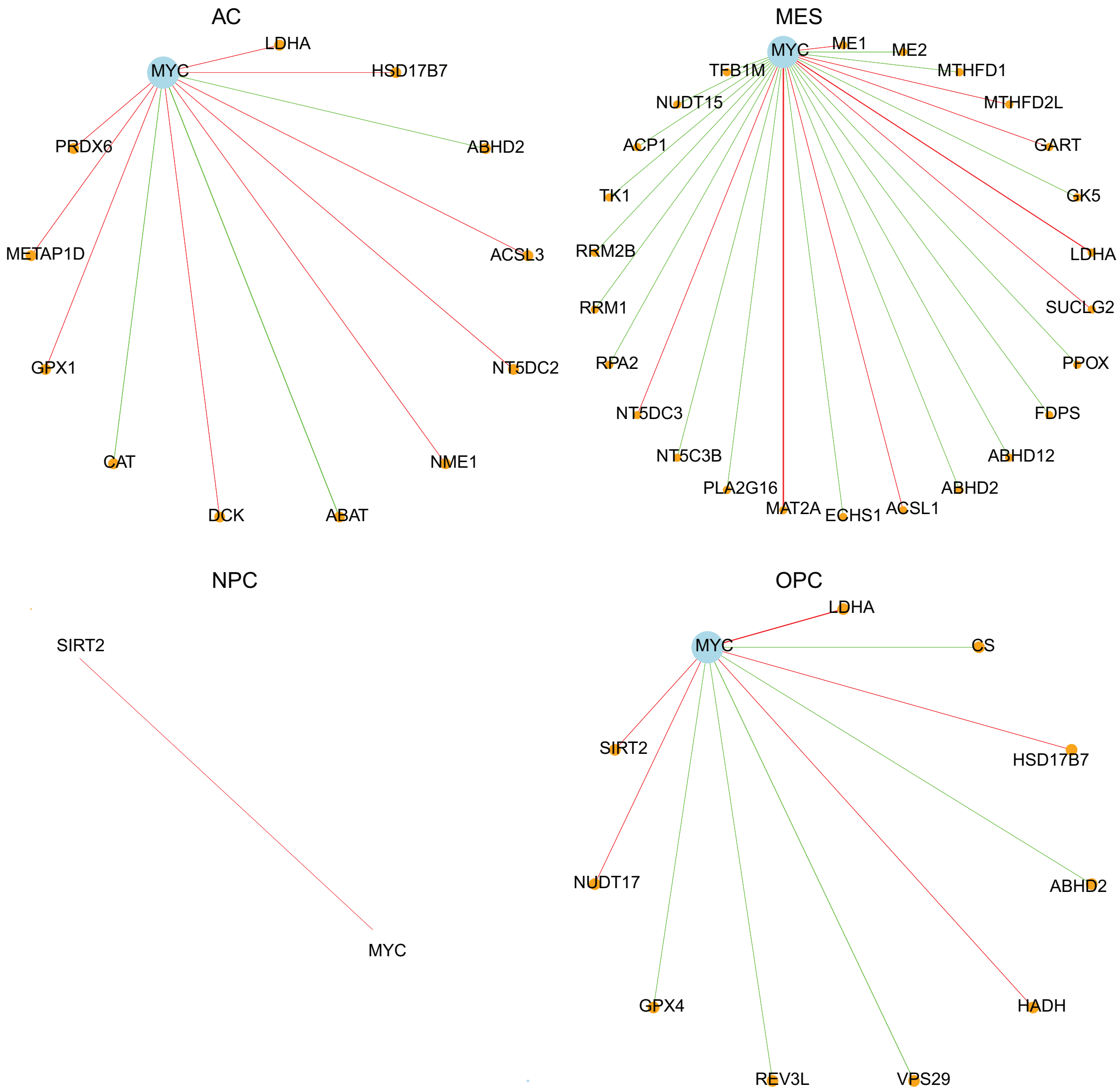


**Figure 14.** Joint network construction results for GBM samples for genes connected to MYC in the metabolism gene set. Connections between non-MYC related genes are not shown.

| pathways | *metabolism enzymes gene set* | Composition of genes connected to **MYC** AC-like | MES-like | OPC-like | Fisher exact test p-value AC | MES | OPC |
|---|---|---|---|---|---|---|---|
| Anaplerotic Reactions of the TCA Cycle | 0.01 | 0 | 0.08 | 0 | 1 | 0.05 | 1 |
| Deamination of Amino Acids: The Urea Cycle | 0.03 | 0 | 0 | 0 | 1 | 1 | 1 |
| Folic Acid Metabolism | 0.03 | 0 | 0.12 | 0 | 1 | 0.02 | 1 |
| Folic Acid Reactions | 0.01 | 0 | 0.04 | 0 | 1 | 0.31 | 1 |
| Gluconeogenesis and Glycolysis | 0.05 | 0.08 | 0.08 | 0.1 | 0.46 | 0.36 | 0.4 |
| Gluconeogenesis and Glycolysis: Regulation | 0.01 | 0 | 0 | 0 | 1 | 1 | 1 |
| Glycolysis and TCA Cycle | 0.05 | 0 | 0.04 | 0.1 | 0.39 | 1 | 0.43 |
| Heme Metabolism | 0.02 | 0 | 0.04 | 0 | 1 | 0.41 | 1 |
| Isoprene Metabolism | 0.07 | 0.08 | 0.04 | 0.1 | 0.59 | 1 | 0.52 |
| Lipid Metabolism | 0.16 | 0.25 | 0.28 | 0.5 | 0.41 | 0.09 | 0.01 |
| Pentose Phosphate Pathways | 0.02 | 0 | 0 | 0 | 1 | 1 | 1 |
| Products of Isoprene Metabolism | 0.08 | 0.08 | 0.04 | 0.1 | 1 | 0.71 | 0.58 |
| Prostaglandins, Thromboxanes, and Leucotrienes | 0.03 | 0 | 0.04 | 0 | 1 | 0.51 | 1 |
| Purine Metabolism | 0.17 | 0.17 | 0.28 | 0.1 | 1 | 0.16 | 1 |
| Pyrimidine Metabolism | 0.09 | 0.25 | 0.16 | 0 | 0.08 | 0.27 | 0.16 |
| Undefined | 0.42 | 0.33 | 0.12 | 0.3 | 0.77 | 0 | 0.53 |
| Total number of genes connected to MYC | (Total: 682) | 12 | 25 | 10 | | | |

**Figure 15.** Gene set enrichment analysis result for genes connected to MYC using the GBM data. To learn functional categories of the genes connected with MYC, these genes are mapped to the NCBI BioSystems database pathways. The number of genes connected to MYC in NPC-like cells is one and hence we do not show NPC-like cells group in this table.
